# A base-modifying enzyme extends the range of anti-phage defense provided by DNA glycosylases

**DOI:** 10.64898/2026.02.25.707755

**Authors:** Adriana Mejía-Pitta, Zhiying Zhang, Amer A. Hossain, Karolina Bartosik, Christian F. Baca, Christopher Peralta, Henrik Molina, Marianna Teplova, Sean F. Brady, Ronald Micura, Dinshaw J. Patel, Luciano A. Marraffini

## Abstract

The most abundant prokaryotic mechanisms of defense against phage predation involve the recognition and destruction of the infecting DNA. One method of counter-defense is the incorporation of modified nucleobases into the phage genome to avoid interaction with enzymes that target the viral DNA. T-even coliphages replace cytosine with 5-hydroxymethylcytosine (5hmC) that in some cases are further decorated with glucosyl groups. To explore the diversity of immunity genes that recognize 5hmC, we infected a library of metagenomic DNA inserts from uncultured, non-sequenced soil bacteria with a mutant T4 phage that harbored only non-glucosylated 5hmC on its genome. Bacteria that resisted infection carried a DNA glycosylase, Brig3, that specifically excises 5hmC nucleobases to generate abasic sites in the phage genome and prevent viral proliferation. The crystal structure of Brig3 bound to its substrate revealed a catalytic mechanism in which the 5hmC nucleobase is flipped out of the DNA into the active site and replaced by an asparagine residue that inserts into the double helix to contact the complementary guanosine. Brig3 is encoded within an operon that also encodes BapA, a hydrolase that removes glucosyl groups from glucosyl-5hmC present in the genome of otherwise Brig3-resistant T-even phages carrying this hypermodified base. Our results uncover a defense strategy in which base-modifying enzymes widen the immune response provided by DNA glycosylases.

**SIGNIFICANCE STATEMENT:** DNA glycosylases provide immunity to bacteria against bacteriophage infection through the excision of modified bases present in the viral genome. Here we describe the discovery of a novel defense DNA glycosylase, Brig3, that excises 5-hydroxymethylcytosine (5hmC) present in the genome of T-even phages. These phages also carry glucosylated 5hmC (glc-5hmC), a hyper-modified base that resist Brig3 activity. We found that Brig3 is assisted by BapA, a glucose hydrolase converts glc-5hmC into 5hmC and generate more Brig3 substrates on the viral genome, increasing the strength of the immune response as well as the range of bacteriophages targetd by Brig3.

## INTRODUCTION

Bacteriophage (phage) infections shape prokaryotic communities by applying constant selective pressure. To survive phage attack, bacteria and archaea have evolved diverse and sophisticated immune systems that target different stages of the phage lifecycle (1, 2). In turn, phages have developed strategies to counteract these defenses, establishing an evolutionary arms race with their hosts (3). A major exponent of the bacteria vs phage genetic contest has been found in the evolution of defense systems that target phage DNA and the different modifications introduced in viral genomes to evade this immunity (4). Restriction-modification and CRISPR-Cas systems are the two most abundant immune systems of prokaryotes (2) and employ sequence-specific nucleases to destroy the incoming phage genome. While restriction enzymes recognize and cleave short DNA sequences, Type I, II and V CRISPR-Cas nuclease complexes are programmed with a guide RNA to find and destroy complementary DNA sequences in the phage genome (5). To counteract these nucleases, some phages introduce nucleobase modifications that prevent the recognition of target DNA sequences in their genomes (6). T-even phages that infect *Escherichia coli* contain 5-hydroxymethylcytosine (5hmC) instead of cytosine (7), which protects them from the nuclease activity of certain restriction enzymes (4). 5hmC nucleotides are incorporated into the phage’s genome during replication and requires several phage-encoded factors. First, deoxycytidine monophosphates nucleotides are hydroxymethylated by Gp42 to generate 5hmC substrates for the replication of the viral DNA (8). Second, to favor the addition of 5hmC into the phage’s genome, (i) cytosines from the nucleotide pool are depleted by the dCTPase Gp56 (9); (ii) the viral (and host) DNA that contains unmodified cytosines is both not transcribed through the activity of the Alc inhibitor (10), and degraded by the DenB endonuclease (11). In response to this mechanism of counter-defense, *E. coli* developed specialized nucleases, encoded by type IV restriction-modification systems, that can cleave phage DNA containing modified bases (12). McrBC (<u>m</u>odified <u>c</u>ytosine <u>r</u>estriction), which cleaves 5-methylcytosine (5mC) and 5hmC-containing DNA, is one of the better characterized type IV restriction enzymes (13). In turn, T-even phages have evolved “hypermodified” nucleobases to protect the phage DNA from cleavage by McrBC (as well as types I, II, and III restriction endonucleases) and the CRISPR nuclease Cas9 (14, 15). T2, T4 and T6 encode glucosyltransferases that introduce glucose groups on 5hmC in the viral DNA (7). In the T4 genome, all 5hmC nucleobases are glucosylated through either α-linkages introduced by α-glucosyl transferase (*a-gt*; 70%) or β-linkages introduced by β-glucosyl transferase (*b-gt*, 30%). T2 lacks *b-gt* but encodes an alternative glucosyltransferase (*g-gt*) that modifies α-glucosyl-5hmC with a second glucosyl group, β-linked to the primary glucose, to generate gentiobiosyl-5hmC (16). As a result of this transferase activity, the T2 genome contains ∼70% of its 5hmC bases α-glucosylated, and ∼5% modified with gentiobiosyl groups (16). T6 also encodes *a-gt* and *g-gt*, but has an inverted proportion of α-glucosyl modifications than T2, with ∼72% of its 5hmC sites doubly glucosylated as gentiobiosyl-5hmC and only ∼3% in the form of α-glucosyl-5hmC (14). Both phages have ∼25% of “naked” 5hmC nucleobases without additional chemical groups. To counter the glucosylation of 5hmC and restore immunity against T-even phages, *E. coli* has evolved type IV restriction-modification systems that specifically recognize them, such as GmrSD, which cleaves glucosylated 5hmC-containing DNA (12).

Through the screening of an environmental DNA (eDNA) library containing large fragments of DNA isolated from an Arizona soil sample, we previously uncovered another strategy used by bacteria to gain immunity against phages carrying modified nucleobases. We identified Brig1 (<u>b</u>acteriophage replication inhibition <u>g</u>lycosylase 1), a DNA glycosylase that, instead of cleaving the phosphate backbone of the phage genome, removes α-glucosyl-5hmC nucleobases to generate abasic sites that prevent viral replication (17). A recent bioinformatic search for homologs of Brig1 resulted in the isolation of Brig2, a DNA glycosylase that provides defense to *Pseudomonas* against infection with phages harboring modified thymine bases (18), a finding that expanded the range of this novel mechanism of immunity. The presence of *brig1* in the soil sample used to make the eDNA library suggested that bacteria from this environment are exposed to phages carrying α-glucosyl-5hmC. Since such phages usually also contain non-glucosylated 5hmC, for example T2 and T6 (7), we reasoned that the eDNA sample could also contain defense systems that target these nucleobases. To test this prediction, we infected the same Arizona soil metagenomic library used to isolate Brig1 with a Δ*a-gt* and Δ*b-gt* double mutant T4 phage that only carries 5hmC on its genome, here abbreviated T4(5hmC). This screen yielded a Brig1 homolog, Brig3, that removes 5hmC nucleobases to generate a basic sites and prevent the replication of T-even phages that harbor unmodified 5hmC nucleobases. X-ray crystallography of Brig3 bound to its DNA substrate revealed that the molecular mechanism of base excision utilizes an asparagine residue (N225) that is inserted in the DNA to replace the 5hmC nucleobase and flip it out of the duplex into the active site. Interestingly, *brig3* is in an operon that also harbors *bapA* (<u>B</u>rig-<u>a</u>ssociated <u>p</u>rotein <u>A</u>), which encodes a glucosyl hydrolase that acts on both α- and β-glucosyl-5hmC residues. BapA converts these hypermodified bases into 5hmC, the substrate for Brig3. Therefore, the combined action of BapA and Brig3 widens the DNA glycosylase immune response to enable defense against T-even phages with genomes that are either partially or completely glucosylated.

## RESULTS

### Brig3 limits the replication of a T4 phage harboring 5hmC nucleobases

To search for defense systems that specifically recognize 5hmC nucleobases, we screened the Arizona soil metagenomic eDNA library (AZ52) that rendered Brig1 (17) with T4(5hmC), an engineered a T4 phage carrying deletions in the *a-gt* and *b-gt* genes. AZ52 contains at least 10 million cosmids harboring a ∼40 kb DNA extracted from soil DNA and transformed into *E. coli* EC100 cells (19) and was infected with T4(5hmC) at a multiplicity of infection (MOI) of 10. Cosmid DNA from surviving colonies was extracted and re-transformed into *E. coli* EC100 cells to obtain a phage-enriched library. Individual transformant colonies were randomly selected to seed top agar media and test for defense against phage T4(5hmC) through the measurement of plaque forming units (PFU). Since phage T5 does not contain modified nucleobases on its genome, it was used as a negative control in this assay. Along with several false positive clones, we found that one of the transformants selected by our screen reduced the PFU values of T4(5hmC), but not T5, phage by at least six orders of magnitude (Fig. S1A).

Sequencing of the cosmid carried by these cells revealed the presence of an eDNA insert containing a *bona fide* bacterial defense island that included a type II-E CBASS system (20) and genes with homology to those found within SspABCD-SspE defense systems (21) (Fig. S1B). This island was not found in any of the deposited sequences in Genbank and the closest sequence identified by nucleotide BLAST of the selected cosmid insert (from the gram-negative, myxobacterial species *Anaeromyxobacter oryzae*) had only 1% of query cover with an E-value of 6E-08. Therefore, we cannot determine the organism that originally harbors the isolated defense island. To test whether the immunity provided by the selected defense island depends on the presence of 5hmC nucleobases, we tested defense against the wild-type phage [T4(wt)] and an engineered phage without cytosine modifications, T4(C) (17). We found that the resistant clone did not affect T4(C) plaquing and reduced T4(wt) PFUs only by one order of magnitude (Fig. 1A). This result suggested that the isolated defense island contains gene/s with potent activity against 5hmC-containing phages and therefore fulfilled the original premise of our eDNA screen.

**Figure 1.**
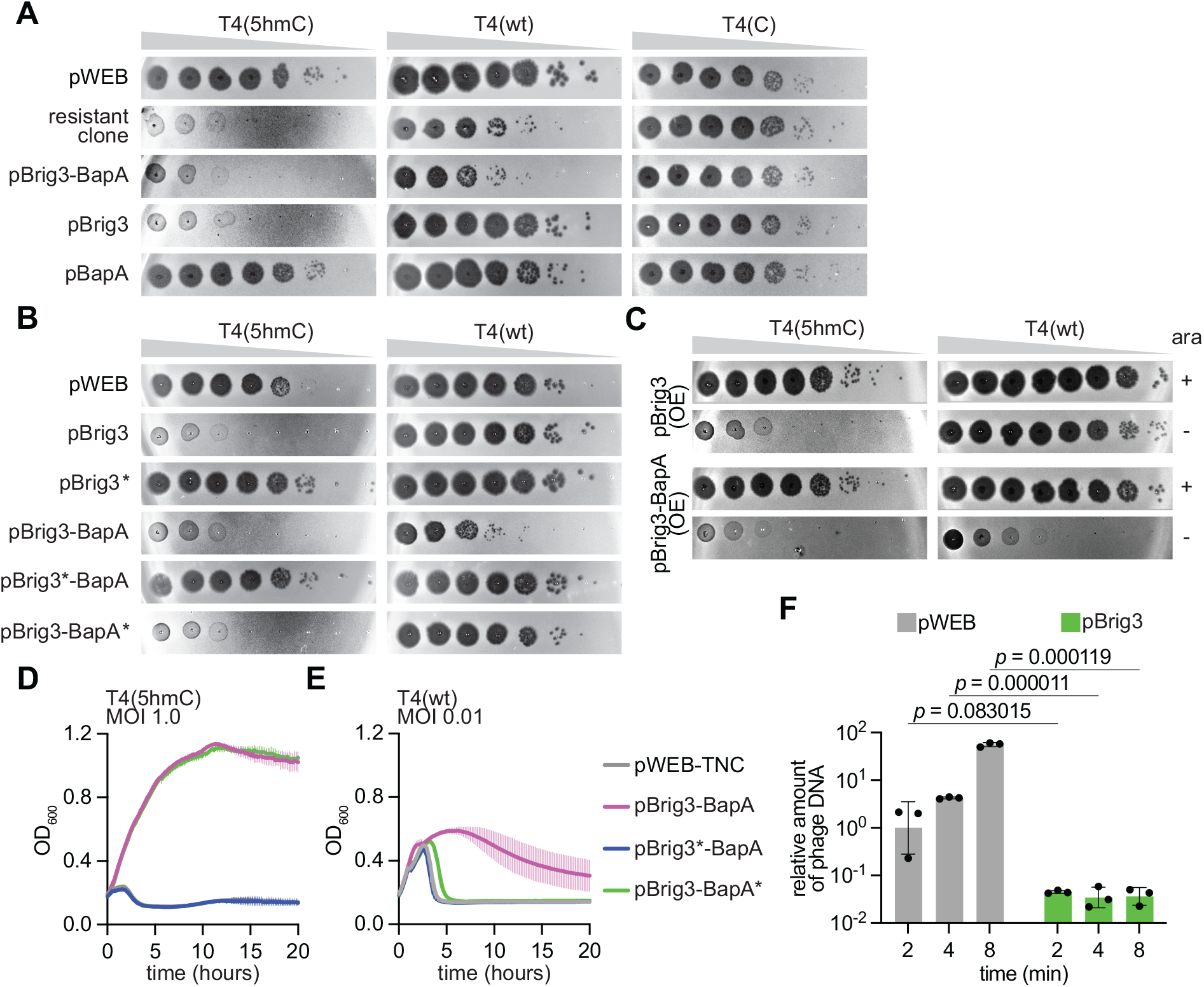
Brig3 provides defense against a T4 phage containing unmodified 5hmC nucleobases. **(A)** Ten-fold serial dilutions of T4 phages containing different cytosine nucleobases on lawns of *E. coli* EC100 that harbor the pWEB cosmid carrying different genes present in the *brig3*-*bapA* operon isolated after the screening of an eDNA library with the T4(5hmC) phage. **(B)** Same as **(A)** but testing the effect of the introduction of stop codons (*) in *brig3* and/or *bapA*. **(C)** Same as **(A)** but testing the effect of the introduction of an arabinose-inducible promoter to drive over-expression (OE) of *brig3* and/or *bapA*; “ara”, arabinose. **(D)** Growth of *E. coli* EC100 that harbor the pWEB cosmid carrying wild-type and stop codon mutant versions of *brig3* and/or *bapA*, measured by optical density at 600 nm (OD_600_) after infection with T4(5hmC) at MOI 1.0. The mean ± S.D. for 3 biological replicates is shown. **(E)** Same as **(D)** but after infection with T4(wt) at MOI 0.01. **(F)** Quantitative PCR analysis of T4(5hmC) DNA through amplification of the *gp34* gene. Viral DNA was extracted from infected *E. coli* EC100 cells carrying pWEB or pBrig3 at 2, 4 and 8 minutes after the addition of phage at an MOI of 1. Fold-change values were calculated relative to the pWEB 2-minute time point. Mean ± SD values are reported for three independent experiments; *p*-values are reported for multiple unpaired Student T-tests.

To identify the genes responsible for immunity, we subcloned six eDNA fragments, F1-F6 (Fig. S1B) containing genes that could be involved in anti-phage defense and re-tested their impact on plaque formation. We found that F6, but none of the other fragments, including F1 which contains the type II-E CBASS system, retained immunity against T4(5hmC) (Fig. S1C). This fragment contained three open reading frames: the first encoding a protein belonging to the superfamily of uracil DNA glycosylases that include Brig1 (17) and Brig2 (18) (Fig. S1D), the second encoding a polypeptide of unknown structure and function, and the third encoding a protein predicted to have ADP-ribosyl glycohydrolase activity (gene predictions were performed with HHpred (22), AlphaFold 2 (23) and DALI (24)). Given that the latter is also present in F2, which did not reduce T4(5hmC) PFUs (Fig. S1C), we discarded its involvement in defense against this phage. The first and second genes of F6 were named *brig3* (<u>b</u>acteriophage <u>r</u>eplication inhibition DNA <u>g</u>lycosylase <u>3</u>, due to its homology to Brig1 and Brig2), and *bapA* (<u>B</u>rig-<u>a</u>ssociated <u>p</u>rotein <u>A</u>). *brig3* and *bapA* overlap with each other and therefore we hypothesized that they are part of an operon which transcription is most likely driven by a promoter located upstream of *brig3* and downstream of the CBASS system, in a region that lacks detectable ORFs (Fig. S1B). We cloned this operon, as well as mutant versions harboring deletions of each gene, into the library cosmid vector pWEB-TNC (25) (abbreviated “pWEB”) and evaluated the different constructs for their ability to decrease T4(5hmC) propagation in plaque assays. We found that *brig3* was solely responsible for defense against T4(5hmC) (Fig. 1A). Interestingly, we could not detect plaque formation, an observation that suggests that mutations in T4(5hmC) that enable escape from *brig3* defense are extremely rare or even lethal for the phage. To test the dependance of *brig3*-mediated immunity on the presence of 5hmC nucleobases, we evaluated the constructs for defense against T4(wt) and T4(C). We observed that *brig3* did not affect the propagation of these phages (Fig. 1A). Interestingly, a reduction in T4(wt) PFUs was detected in the presence of both *brig3* and *bapA* (also observed with the cosmid harboring the full defense island) (Fig. 1A). These results were corroborated by eliminating translation via the introduction of stop codons instead of deletions (Fig. 1B), and therefore we conclude that proteins, rather than RNA molecules, are responsible for defense. Similar results were obtained after measuring cell survival (following the optical density at 600 nm, OD_600_, of *E. coli* cultures) upon infection, instead of PFU. *brig3* was sufficient to protect bacteria after addition of T4(5hmC) at a multiplicity of infection (MOI) of 1 (Fig. 1D) and 10 (Figs. S2A-B), but not against infections with T4(wt), even at low MOIs of 0.01, 0.1 nor 1.0 (Figs. 1E, S2C-D).

Next, we investigated the effects of different levels of Brig3 expression on defense. First we integrated a copy of *brig3* and its upstream sequence (presumably containing a promoter) into the *attTn7* site of *E. coli* EC100, downstream of the *glmS* gene (26), generating the strain gBrig3. In order to compare the plasmid (multi-copy)-and chromosome (single copy)-derived levels of Brig3 expression, we added a hexa-histidyl tag at the C-terminus of the protein, generating plasmid pBrig3-His_6_ and strain gBrig3-His_6_, and performed western blots using recombinant Brig3-His_6_ (see below) and a lysate of cells harboring the pWEB-TNC vector as positive and negative controls, respectively. As expected, we detected strong plasmid-borne expression of Brig3 but a much lower signal in the presence of a single genomic copy of the gene (Fig. S2E). We then compared the immunity against T4(5hmC) phage. We found that the ability of Brig3 to reduce plaque formation (Fig. S2F) and enable culture growth (Fig. S2G) upon infection was affected by the addition of the hexa-histidyl tag, especially when expressed from a single chromosomal copy. Regarding the effect of Brig3 expression on defense, we found that gBrig3 provided strong defense against T4(5hmC) infection, but at lower levels than the highly abundant pBrig3 copy (Figs. S2F-G). We also investigated the effect of *brig3* over-expression, specifically to determine if higher expression could mediate defense against T4(wt) phage, by cloning it on a plasmid under an arabinose-inducible promoter (P*_ara_*). While T4(5hmC) PFUs were reduced only upon the addition of arabinose, Brig3 over-expression was not sufficient to preventT4(wt) propagation (Fig. 1C). Likewise, we observed inducer-dependent culture growth after infection with T4(5hmC) at MOIs 1.0 and 10 (Figs. S2H-I) but not after infection with T4(wt) at MOIs 0.1 and 1.0 (Figs. S2J-K).

Finally, we performed quantitative PCR (qPCR) of the T4 *gp43* and *gp34* loci using DNA extracted from *E. coli* harboring Brig3 (or a vector control) at 2, 4 and 8 minutes after infection with T4(5hmC). While defenseless cells showed high levels of DNA accumulation, Brig3 expression led to significantly lower levels of viral DNA (Figs. 1F and S2L), a result that demonstrates that Brig3 limits T4(5hmC) DNA replication.

Altogether, these data demonstrate that different levels of the putative DNA glycosylase Brig3, can prevent the propagation of T4(5hmC), but not of wild-type T4 phage, likely by affecting viral DNA replication.

### Brig3 provides immunity against diverse phages harboring 5-hydroxymethylcytosine nucleobases

We also tested immunity against other T-even phages with genomes that contain unmodified 5hmC residues, T2 and T6. Similarly to the results obtained for T4(5hmC), Brig3 prevented plaque formation (Fig. 2A). Infection of liquid cultures with each of these phages produced equivalent results, showing bacterial growth at both MOI 1 (Figs. 2B-C) and 10 (Fig. S3A-B). In contrast to the T4(5hmC) genome, in which all guanines are paired with unmodified 5hmC bases, T2 and T6 phages decorate ∼75% of their 5hmC residues with α-glucosyl modifications, i.e. only ∼25% of the 5hmC bases are unmodified (14). We conclude from these data that Brig3 can restrict infection by phages that have a small fraction of 5hmC in their genomes. We further tested this by infecting cells carrying pBrig3 with T4 mutant phages that express only one of the two glycosyltransferases, T4(Δ*b-gt*) and T4(Δ*a-gt*), which carry only α- and β-glucosyl-5hmC bases in their genomes, respectively. Because β-glucosyltransferase can act on adjacent 5hmC substrates, the T4(Δ*a-gt*) genome is completely glucosylated and lacks unmodified 5hmC nucleobases required for Brig3 activity (7, 27). In contrast, α-glucosyltransferase only modifies one of two adjacent 5hmC residues (7, 27), leaving ∼25% of 5hmC unmodified nucleobases in the T4(Δ*b-gt*) genome, similarly to T2 and T6. We found that Brig3 strongly prevented T4(Δ*b-gt*) plaque formation but had a more limited impact on T4(Δ*a-gt*) propagation, reducing PFU by less than 10-fold, even when over-expressed using an arabinose-inducible promoter (Fig. S3C). Therefore, these results corroborate that Brig3 can provide immunity to phages that carry a limited number of unmodified 5hmC nucleobases in their genomes. Finally, we tested Brig3 immunity against eleven T-even phages harboring 5hmC nucleobases (Bas35 to Bas45) from the BASEL phage collection (28) and found a marked reduction in plaque formation in all infections (Fig. 2D). In contrast, plaquing by phages from this collection that do not belong to the *Tevenvirinae* family was not affected by Brig3 (Fig. S3D).

**Figure 2.**
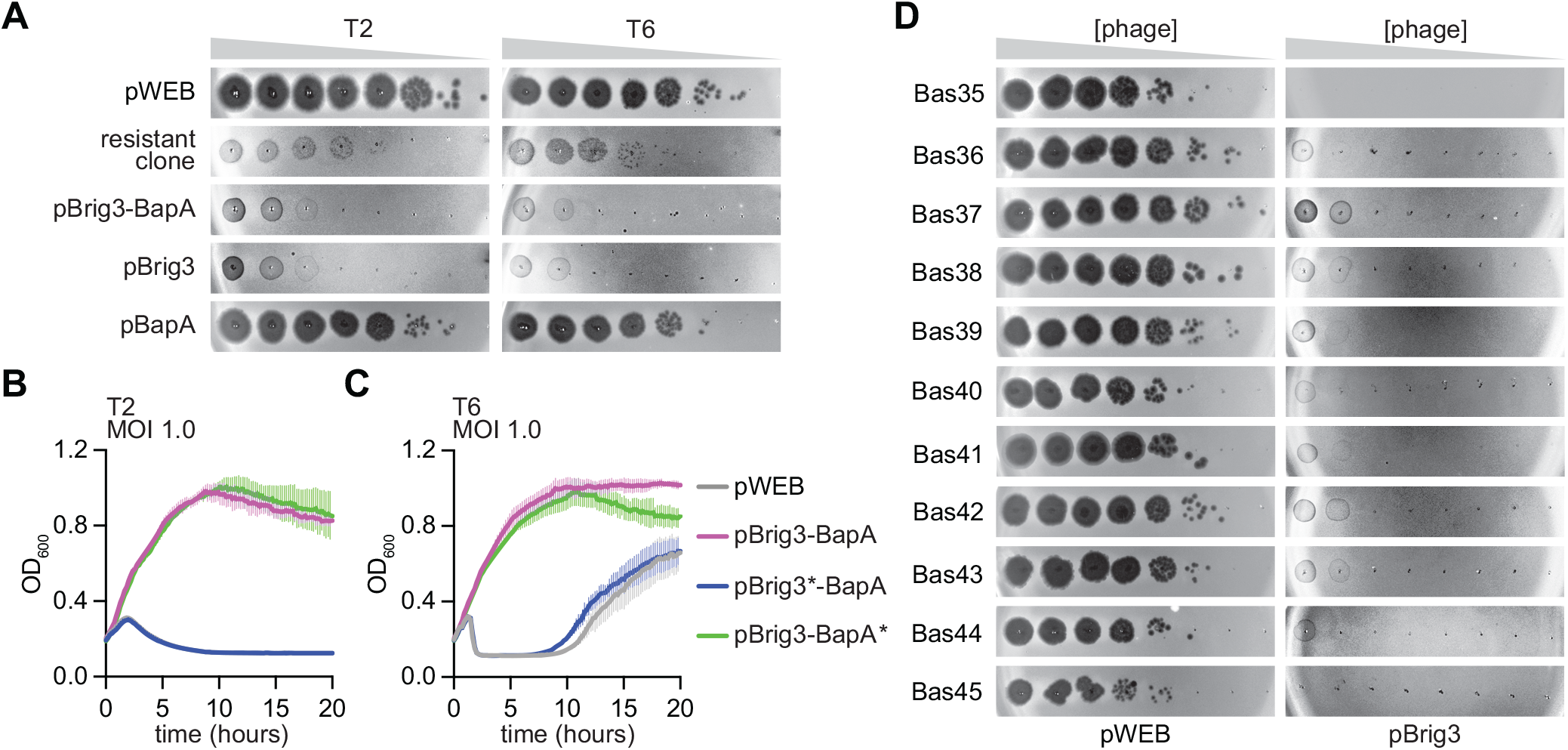
Brig3 provides immunity against diverse T-even phages. **(A)** Ten-fold serial dilutions of T2 or T6 phages on lawns of *E. coli* EC100 that harbor the pWEB cosmid expressing Brig3 and/or BapA. **(B)** Growth of *E. coli* EC100 that harbor the pWEB cosmid carrying wild-type and stop codon mutant versions of *brig3* and/or *bapA*, measured by optical density at 600 nm (OD_600_) after infection with T2 at MOI 1.0. The mean ± S.D. for 3 biological replicates is shown. **(C)** Same as **(B)** but after infection with T6. **(D)** Ten-fold serial dilutions of phages from the BASEL collection Bas35-45 spotted on lawns of *E. coli* EC100 carrying pWEB or pBrig3.

Altogether these data demonstrate that Brig3 provides strong defense against diverse phages carrying 5hmC nucleobases in a fraction of their genomes.

### Brig3 excises 5hmC nucleobases to generate abasic sites

Given that Brig3 is a homolog of the DNA glycosylase Brig1 (Fig. S1D) and that it provides immunity against phages that contain 5hmC nucleobases in their genomes (Fig. 1A), we hypothesized that the mechanism of defense involves the excision of these bases from the viral DNA. To test this, we purified a C-terminal hexa-histidyl tagged version of Brig3 (Fig. S4A-B) and incubated it with a 60-nt DNA oligonucleotide (Fig. 3A) containing in position 21 a single 5hmC, 5-methylcytosine (5mC) or cytosine (C) nucleobase (reactions contained 1 µM Brig3 and 1 µM of oligonucleotide). Due to the weak homology of Brig3 to uracil DNA glycosylases, we also tested activity on oligonucleotides harboring uracil (U) instead of 5hmC, or thymine (T) as control (Fig. 3A). After treatment of these different ssDNA substrates with purified Brig3, we detected base excision upon heating the reactions to 65°C in the presence of NaOH, which catalyzes a β-elimination reaction at the abasic site that results in cleavage of the phosphate bond within the oligonucleotide substrate (29). We found that Brig3 excises both 5hmC and U, but not C, 5mC nor T, bases (Fig. 3B). As a control we treated the oligonucleotides with a commercially available human uracil DNA glycosylase (hSMUG1), which generated abasic sites only on U-containing oligonucleotides (Fig. 3B). To compare the activities of Brig3 on 5hmC and U substrates, we incubated the oligonucleotides harboring these nucleobases with an increasing concentration of enzyme. We found that while uracil excision required at least 1 µM of enzyme (Fig. 3C), activity against 5hmC nucleobases was achieved using Brig3 at a concentration of 10 nM (Figs. 3C and S4C). This result indicates that although Brig3 displays a residual activity against uracil, its main substrate is 5hmC. We also corroborated the activity of Brig3 on 5hmC-containing substrates with a different method that detects the presence of abasic sites on DNA. We treated Brig3 reaction products with Endonuclease IV, an apurinic/apyrimidinic (AP) endonuclease that cleaves the DNA sugar-phosphate backbone adjacent to an abasic site (30) and observed cleavage of the ssDNA oligonucleotide into two fragments of ∼20 and ∼40 nt (Fig. S4D), most likely at position 21 where the 5hmC nucleobase is located (Fig. 3A), further demonstrating base excision by Brig3. Finally, to unequivocally determine that Brig3 causes 5hmC excision, we analyzed 18-nt ssDNA oligonucleotides containing in position 9 either cytosine, 5hmC, α-glucosyl-5hmC or β-glucosyl-5hmC (Fig. S4E) by liquid chromatography and high resolution/high mass accuracy mass spectrometry. Glucosylated substrates were obtained by treatment of the 5hmC-containing oligonucleotide with either α-or β- glucosyltransferase. In the former case, the low efficiency of the enzymatic reaction prevented full conversion of the 5hmC nucleobases to α-glucosyl-5hmC (Fig. S4F).

**Figure 3.**
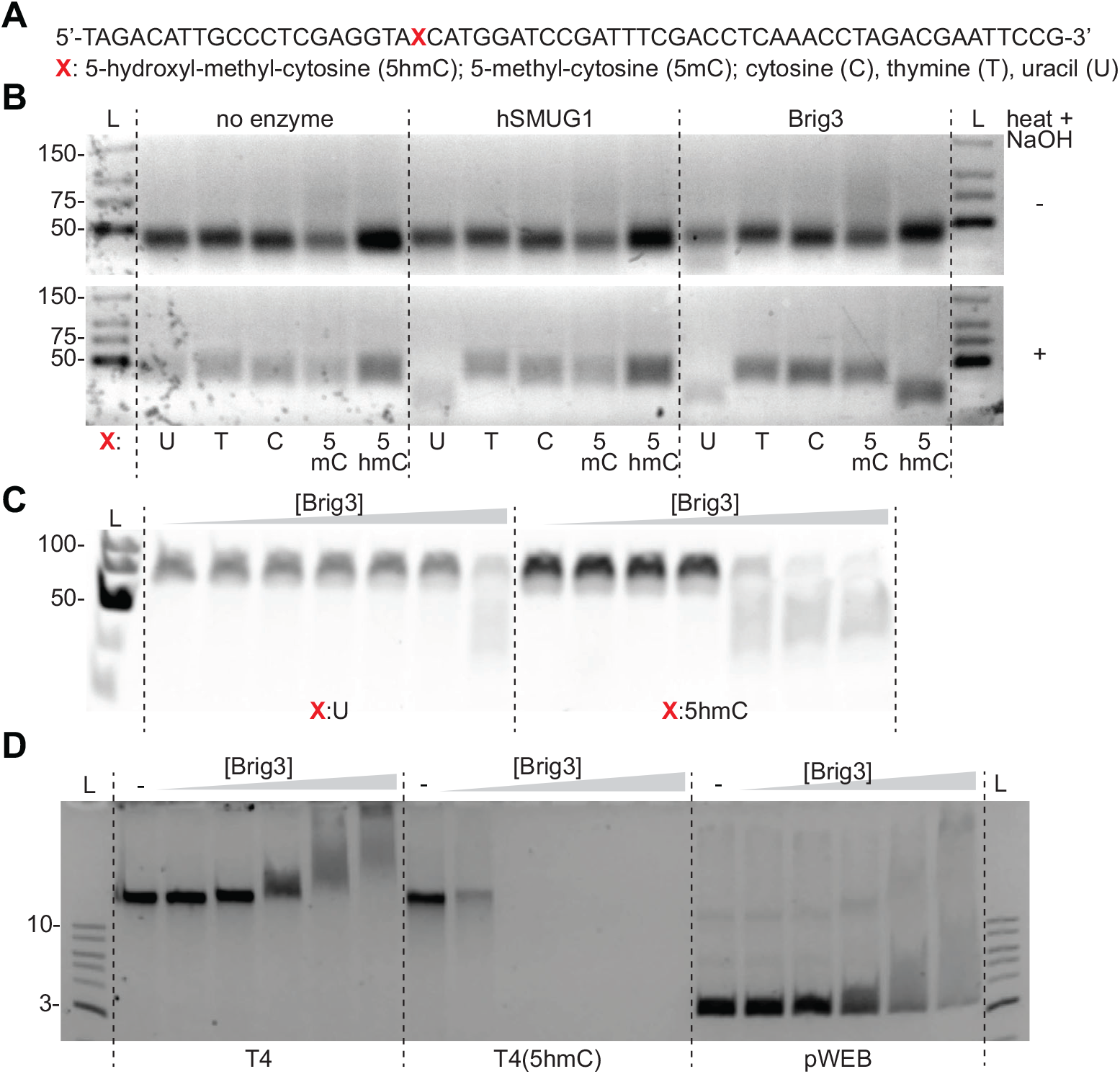
Brig3 excises 5hmC nucleobases. **(A)** Sequence of the 60-nt single-stranded oligonucleotide used to test Brig3 activity. The position 21 from the 5’ end (red “X”) is occupied by different nucleobases. **(B)** PAGE of different oligonucleotides either untreated or incubated with hSMUG1 or Brig3 at 37°C overnight, with and without heating in the presence of NaOH for 30 minutes. Gels were stained with ethidium bromide. L, ssDNA size ladder. **(C)** Same as **(B)** but incubating U-or 5hmC-containing oligonucleotides with ten-fold dilutions of Brig3 (10^-3^, 10^-2^, 10^-1^, 1, 10, 100, 1000 nM). **(D)** Agarose gel electrophoresis of T4, T4(5hmC) or pWEB DNA (12.5 ng) treated with increasing concentrations (2, 20, 200, 400, 800 nM) of Brig3 for 30 minutes at 37°C. L, DNA size ladder.

Upon treatment with Brig3, only the signal for the 5hmC substrate shifted to a lower molecular weight that corresponded to the presence of an AP site, including the signal of the unmodified 5hmC substrate present after treatment with α-glucosyl-transferase (Fig. S4F). Finally, we tested the activity of Brig3 on dsDNA substrates by pairing the 60-nt ssDNA oligonucleotides containing either 5hmC or cytosine with their complementary counterpart, the former also containing 5hmC (Fig. S4G). We incubated these substrates, along with the ssDNA substrates as controls, with Brig3 and heated the reactions to 65°C in the presence of NaOH to trigger DNA cleavage at abasic sites by β-elimination. PAGE of the reaction products showed that only the dsDNA substrate containing the 5hmC modifications was fragmented (Fig. S4H), a result that demonstrates Brig3 base excision activity on modified dsDNA, the DNA form present in the genome of T-even phages.

Next, we tested the effect of purified Brig3 on phage DNA. We treated viral DNA extracted from T4(wt) and T4(5hmC) phages, as well as plasmid DNA propagated in an *E. coli* strain devoid of DNA modification enzymes, with increasing concentrations of Brig3 for 30 minutes at 37°C. Agarose gel electrophoresis of the reaction products revealed full degradation of the T4(5hmC) DNA, even at very low concentrations of the Brig3 enzyme (Fig. 3D). Degradation is most likely a result of β-elimination after generation of multiple abasic sites in the phage genome, due to the high temperature reached during electrophoresis as observed previously for T4(wt) DNA treated with Brig1 (17). In contrast, T4(wt) and plasmid DNA remained intact and displayed a mobility shift that indicates binding of Brig3 without base excision (Fig. 3D). This observation aligns with the hypothesis that uracil DNA glycosylases bind to non-target DNA to search for their substrates (31). Altogether, these results demonstrate that Brig3 is a 5hmC DNA glycosylase, and suggest that it evolved divergently from canonical members of the uracil DNA glycosylase superfamily to provide phage defense.

### An asparagine residue displaces 5hmC into the glycosylase active site to promote base excision

To understand the interactions of Brig3 with 5hmC-modified DNA, we purified Brig3 harboring a C-terminal hexa-histidyl tag using Ni-NTA affinity chromatography followed by size exclusion chromatography (Fig. S4A) and used X-ray crystallography to obtain the structure of this DNA glycosylase with and without its substrate. Apo-Brig3 structure (solved at 3.0 Å resolution; statistics listed in Table S1) is a single-domain protein composed of a mixed α/β topology consisting of nine α helices and six β strands (Fig. S5A). The core is formed by a curved, mixed β sheet (Fig. S5A), similar to the human alkyladenine glycosylase AAG (32) (Fig. S5B), as well as other previously reported for DNA glycosylases (32–34), both in terms of overall fold and catalytic pocket architecture (Figs. S5C-D).

We also solved the co-crystal structure of Brig3 bound to a 12-mer dsDNA duplex containing a single central 5hmC modification (Fig. S6A) at 1.7 Å resolution (statistics listed in Table S1). The electron density map provided sufficient detail to accurately refine the complex and trace both the protein and the dsDNA substrate (Fig. 4A). The structure revealed the absence of 5hmC density, which indicated the occurrence of base excision within the crystals (Fig. 4B). The AP site displayed the sugar ring inserted into the DNA duplex with intact 5’ and 3’ phosphodiester bonds (Fig. 4C). These findings further demonstrate that Brig3 acts as a 5hmC DNA glycosylase without DNA lyase activity, as is the case for some DNA glycosylases involved in DNA repair (35).

**Figure 4.**
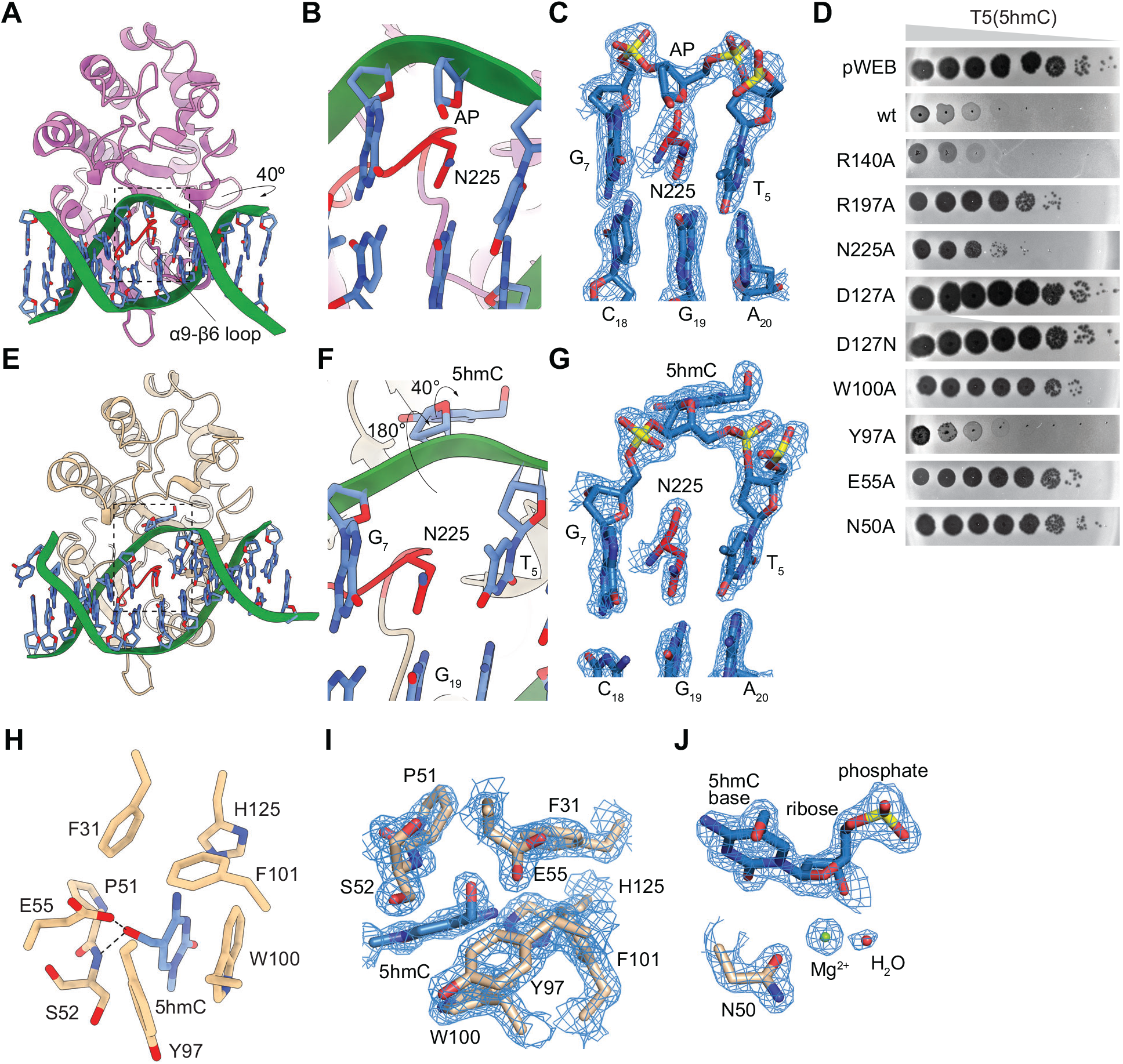
Crystal structure of Brig3 bound to 5hmC-containing DNA. **(A)** Wild-type Brig3 (pink) bound to a 12-mer dsDNA duplex (green; sequence is shown in Fig. S6A). The dsDNA originally contained a central 5hmC nucleobase, which was excised to generate an inward-directed abasic site. **(B)** Region highlighted by the inset in **(A)**. Residue N225 is inserted between the base pairs flanking the abasic site (apurinic/apyrimidinic site, AP). **(C)** Electron density map showing the inward-directed abasic site and the R225 residue inserted between G_7_-C_18_ and T_5_-A_20_ base pairs (see Fig. S6A for numbering), 2Fo-Fc=2.0 σ. **(D)** Ten-fold serial dilutions of T4(5hmC) phage spotted on lawns of *E. coli* EC100 carrying pWEB or pBrig3, encoding either the wild-type protein or mutant versions carrying different amino acid substitutions. **(E)** Brig3^D127N^ (gold) bound to a 12-mer dsDNA duplex (green; sequence is shown in Fig. S7G) containing a central 5hmC nucleobase, whose base and sugar was flipped out of the duple into the active site. **(F)** Region highlighted by the inset in **(E)**. Residue N225 is inserted between the base pairs flanking the flipped 5hmC nucleobase, with a rotation of the ribose ring of approximately 180° and of the nucleobase of about 40° clockwise around the glycosidic bond. **(G)** Electron density map showing the flipped 5hmC nucleobase and the R225 residue inserted between G_7_-C_18_ and T_5_-A_20_ base pairs (see Fig. S7G for numbering), 2Fo-Fc=2.0 σ. **(H)** Brig3 catalytic pocket showing the flipped 5hmC nucleobase surrounded by aromatic residues. **(I)** Electron density map of Brig3 aromatic-lined catalytic pocket occupied by the flipped 5hmC nucleobase, 2Fo-Fc=2.0 σ. **(J)** Electron density map showing a divalent cation (represented as Mg²⁺) near the glycosidic bond of 5hmC, which, along with an adjacent water molecule and the side chain of N50, could play a crucial role in the base excision reaction, 2Fo-Fc=2.0 σ.

The bound DNA adopts a B-DNA type helix that is trapped within the positively charged protein core (Fig. S6B), with an overall bend of 40° away from Brig3 (Fig. 4A). R140, located in the loop region between α7 and β2, and R197, situated in the hairpin loop between β4 and β5, interact with the phosphodiester backbone on opposing strands of the DNA (Figs. S6B-C). Brig3 DNA-binding surface is relatively flat with the exception of the long loop between α9 and β6, which protrudes from the binding surface and inserts into the DNA double helix to occupy the place of the 5hmC nucleobase (Fig. 4A), likely after the occurrence of the base excision reaction. N225 within this loop replaces the 5hmC base, inserting between the adjacent base pairs and inducing a slight distortion in the DNA (Fig. 4B-C, with ideal electron density in this region). N225 forms a hydrogen-pi electron bond with the neighboring base G_7_ and interacts with N1 and C2-NH_2_ of the opposite and complementary base G_19_, while also forming hydrogen bonds with the O4’ atom of the sugar ring at the AP site, thereby stabilizing the distorted DNA structure (Fig. S6D). The remainder of the DNA-binding interface is formed by an alpha helix and a connecting loop, with A174, E175, G229, T230, and N233 mediating interactions with the DNA via the main chain (Figs. S6C, E). Finally, we did not observe significant conformational changes generated by the binding of 5hmC-dsDNA, with a root mean square deviation (RMSD) of 0.36 Å between the apo and DNA-bound Brig3 structures (Fig. S6F). The only notable shift occurred in the α9-β6 loop that harbors N225, which moves closer to the minor groove upon DNA binding (Fig. S6G). To test our structural findings, we made alanine substitutions of the key amino acids and tested their impact on phage defense. We introduced the R140A, R197A or N225A mutations into Brig3 and infected the different strains with T4(5hmC). We found that while the R140A mutation did not impair Brig3 activity, R197A and N225A substitutions disrupted defense and led to an increase in PFUs of six and five orders of magnitude, respectively (Fig. 4D), a result that corroborates the importance of these two residues for Brig3 activity.

To better understand the molecular details of 5hmC recognition and excision, we decided to obtain the co-crystal structure of a catalytically inactive Brig3 mutant in complex with a DNA substrate. We wanted to make a substitution that would not cause major structural distortions in the active site and therefore enable trapping the substrate in a pre-active state. We inspected the Brig3:DNA structure and found one such residue, D127, pointing towards an inner cavity adjacent to the AP site that most likely constitutes the binding pocket for the 5hmC nucleobase (Fig. S7A). Importantly, D127 does not make direct contact with the phosphate backbone (Figs. S6C, S6E and S7A) and therefore its mutation should not drastically disrupt DNA binding. We tested whether alanine and asparagine substitutions of D127 affected Brig3 defense using a plaquing assay and found that both mutations prevent immunity against T4(5hmC) (Fig. 4D). We also purified C-terminal hexa-histidyl-tagged versions of the mutant proteins (Figs. S7B-E) and use them to treated 60-nt DNA oligonucleotide substrates containing a single 5hmC nucleobase, detecting base excision by heating the reactions to 65 °C in the presence of NaOH before separation of the products by polyacrylamide gel electrophoresis (PAGE). We found that both mutations prevented cleavage of the oligonucleotide (Fig. S7F), a result that corroborates the importance of D127 for Brig3 activity. We next solved the co-crystal structure of Brig3^D127N^ and a 12-mer DNA duplex. We used the D127N mutant because asparagine maintains a similar size and orientation as aspartic acid but cannot act as a general base. Since we were unable to obtain high quality crystals with the DNA duplex used for co-crystallization of wild-type Brig3 (Fig. S6A), we used a substrate with a 5’-G overhang on the strand harboring the 5hmC base (Fig. S7G). The electron density map provided sufficient detail to accurately refine the complex and trace both the protein and the bound DNA at a 1.6 Å resolution (statistics listed in Table S1). The structure revealed that both the base and sugar of the 5hmC nucleoside are flipped out of the DNA duplex (Figs. 4E-G) into an aromatic-lined catalytic pocket (Figs. 4H-I). Within this pocket, the 5hmC base is stacked by F31, W100, F101 and H125, with Y97 and the peptide bond of P51 also participating in the binding (Figs. 4H-I). These side chains adopt the same orientation as in the wild-type Brig3-DNA complex, suggesting that the shape of the pocket is predetermined rather than being induced by substrate binding (Fig. S7H). While aromatic cage capture positions 5hmC for catalysis, hydrogen bonds within this pocket account for the selectivity of Brig3 for this base. Specifically, E55, as well as the nitrogen atom of the peptide bond of S52, form a hydrogen bond with the oxygen of the CH_2_OH group of 5hmC base (Fig. 4H), providing a key structural explanation for the recognition of this base over 5-methylcytosine or cytosine. Regarding catalysis of the base excision reaction, we identified a divalent cation (assumed to be Mg^2+^) bridging the side chains of N50 and the C1’ of 5hmC, in the vicinity of the glycosidic bond to be cleaved (Fig. 4J). It is conceivable that the Mg^2+^ and the nearby water molecule play an important role in the hydrolysis of the 5hmC base to generate the AP site. We tested these structural findings in vivo to evaluate the importance of the key residues in Brig3-mediated defense. We made alanine substitutions of W100, Y97, E55 and N50 and infected lawns of bacteria expressing the different mutants with T4(5hmC). We found that W100A, E55A and N50A mutations completely abrogated Brig3 defense (Fig. 4D). In contrast, Y97A displayed a minor effect on immunity, increasing PFUs by only two orders of magnitude (Fig. 4D). These data confirm the contribution of these residues for Brig3 catalysis.

Similarly to the aromatic cage described above (Fig. S7H), the overall conformations of wild-type and D127N Brig3:DNA complexes display minimal differences, with an RMSD of 0.40 Å (Fig. S7I). A slight change occurs in the positioning of the insertion loop between α9 and β6, with a displacement of less than 2 Å (Fig. S7J). In the D127N mutant, the side chain of N225 that projects from this loop displays the same interactions with the adjacent G7 base and opposite G19 base as the wild-type enzyme (Figs. 4E-G and S7K). At this site, however, while in wild-type Brig3 the baseless sugar ring remains within the DNA duplex (Figs. 4A-C), in the D127N mutant it rotates out of the double helix approximately by 180° and the 5hmC nucleobase rotates clockwise by about 40° around the glycosidic bond (Figs. 4F). This rotation of the glycosidic bond exposes the Watson-Crick edge of the base to the deep pocket next to the enzyme active site (Fig. 4H). As a result of the flipping of the 5hmC nucleoside, the phosphate backbone at its 5’ end suffers a 66° bend, when compared to the DNA structure within wild-type Brig3, that causes a shift of the double helix of ∼7 Å towards the mutant protein (Fig. S7I). To maintain the double helix, the opposite strand shifts ∼6 Å in the same direction (Fig. S7I), allowing the side chains of R53 and Y97, as well as the peptide bonds of S139, A174, E175 and A224, to interact with its DNA phosphate backbone and expand the atomic connections between Brig3 and its substrate (Figs. S7L-M). In contrast, the phosphate backbone at the 3’ end of the 5hmC nucleoside adopts the same conformation as in wild-type Brig3, engaging in similar hydrogen bond interactions with the amino group of R197 and the peptide bonds of G229, T230 and N233 (Figs. S7L-M). In summary, our structural analysis reveals the molecular details of Brig3 activity, revealing a mechanism in which binding of the DNA substrate results in the flipping of the 5hmC nucleoside into an active site pocket lined with aromatic residues to catalyze base excision.

### BapA removes glucose residues from α- and β-glucosyl-5hmC to expand Brig3 immunity

As mentioned above, in addition to drastically preventing T4(5hmC) plaque formation, we noticed that the library clone selected in our screen provided partial protection against T4(wt), reducing PFUs by two orders of magnitude (Fig. 1A). Further fragmentation and introduction of stop codons showed that, in addition to Brig3, *ORF2* (*bapA*, see above) is required for defense against T4(wt) (Figs. 1A-B). We noticed that the immunity provided by Brig3 and BapA against T4(wt) is weaker than that provided by Brig3 alone against T4(5hmC). This was the case both when we measured PFU generation (Figs. 1A-B) and in cell survival assays, which showed that Brig3 and BapA were required to maintain a temporary growth of cultures infected with T4(wt) at an MOI of 0.01 and 0.1, but protection was minimal at MOI 1 (Figs. 1E and S2C-D), compared to the strong growth observed after treatment of cultures producing Brig3 with T4(5hmC) at high MOIs, 1 and 10 (Figs. 1D and S2B). We wondered whether this weaker defense was a consequence of low expression of *brig3*-*bapA* and therefore cloned the operon under the control of an arabinose-inducible promoter to increase its transcription. Indeed, over-expression resulted in improved protection of *E. coli* from T4(wt), both by further reducing PFUs to the same levels observed after plaquing of T4(5hmC) on lawns of bacteria expressing Brig3 alone (Fig. 1C), as well as by enabling more robust growth of infected cultures at MOIs 0.1 and 1.0 (Figs. S2G-H). In contrast to the results for T4(wt), the combined presence of Brig3 and BapA did not increase defense against T2 and T6 infection, which genomes have ∼25% of 5hmC unmodified nucleobases, measured by both phage plaquing (Fig. 2A) and by the survival of infected cultures (Figs. 2B-C and S3A-B).

Given the specificity of Brig3 for 5hmC nucleobases, we hypothesized that the requirement of BapA to provide defense against T4(wt) infection, a phage in which all 5hmC bases are glucosylated, may be due to an activity during viral replication that either (i) inhibits glucosyltransferases, (ii) removes glucose residues from the DNA, or (iii) expands Brig3 substrate specificity to α- or β-glucosyl-5hmC. To explore these possibilities, we first expressed a N-terminal hexa-histidyl-SUMO tagged version of BapA to purify a tagless version of the protein (Fig. S8A-B) and incubated it with purified Brig3-His_6_ to allow for complex formation before protein separation using size exclusion chromatography (Figs. S8C-D). Since we did not observe co-purification of the two proteins, we therefore conclude that they do not form a stable complex. Next, we performed in vitro base excision assays with the purified proteins and the 60-nt 5hmC-containing oligonucleotide substrate used in previous experiments (Fig. 3A), treated with purified α- or β-glucosyltransferase to glucosylate the 5hmC base (17) (Fig. 5A).

**Figure 5.**
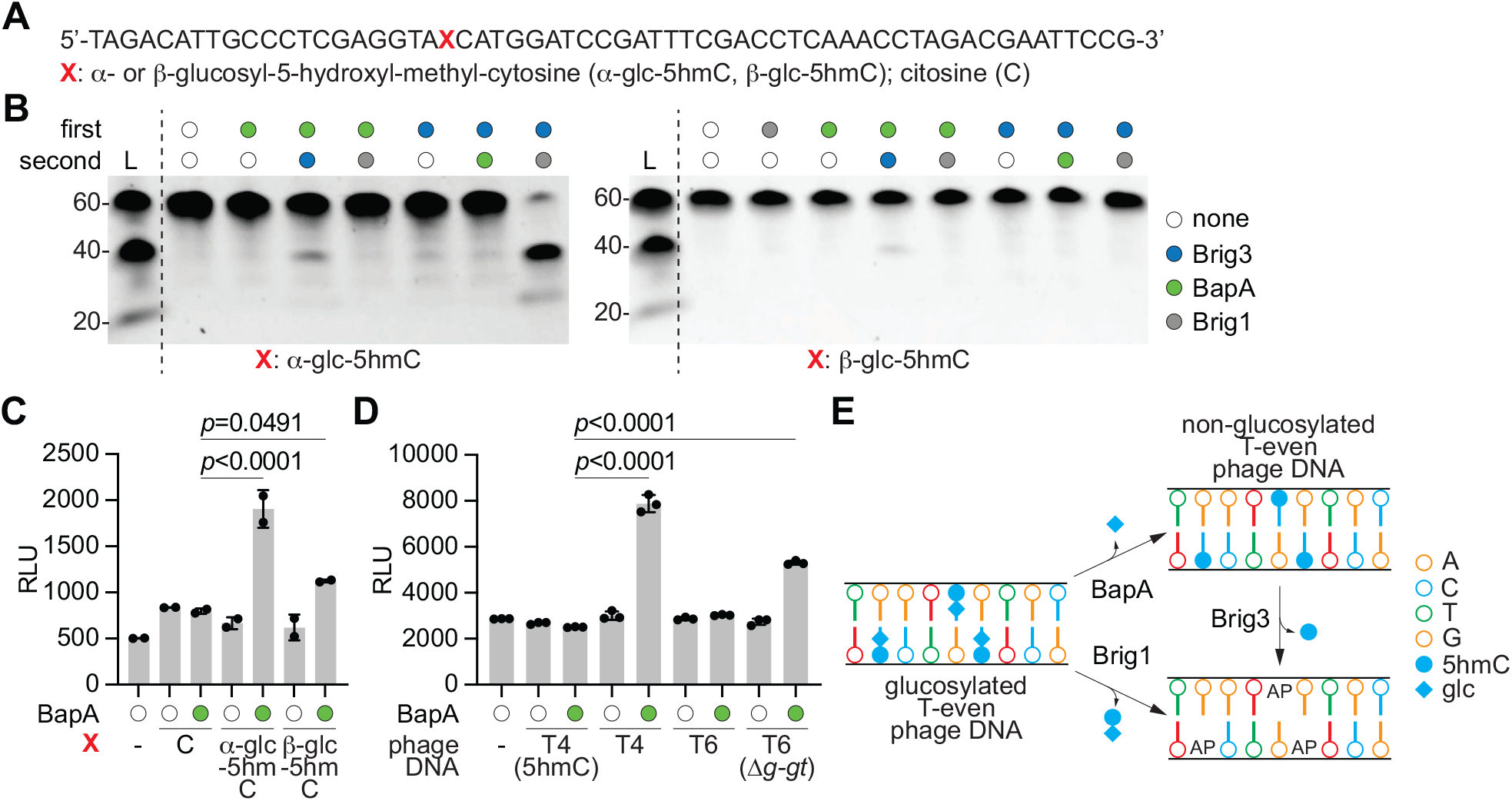
BapA releases glucose from α- and β-glucosyl-5hmC nucleobases. **(A)** Sequence of the 60-nt single-stranded oligonucleotide used to test Brig3/BapA activity. The position 21 from the 5’ end (red “X”) is occupied by different nucleobases. **(B)** PAGE of α-(left gel) or β-(right gel) glucosyl-5hmC oligonucleotides either untreated or incubated at 37°C overnight with the first enzyme and then at 37°C for one hour with the second enzyme, followed by heating in the presence of NaOH for 30 minutes prior to electrophoresis. Gels were stained with ethidium bromide. L, ssDNA size ladder. **(C)** Luminescence assay for glucose detection after incubation of oligonucleotides shown in **(A)** with BapA overnight at 37°C. RLU, relative luminescence units. Error bars represent the standard error of the mean; *p*-values are reported for a Ordinary One-Way ANOVA with multiple comparisons. **(D)** same as **(C)** but treating genomic DNA extracted from different phages with BapA. **(E)** Model comparing Brig1 and Brig3/BapA immunity. Brig1 and BapA act on the genomes of T-even phages that contain glucosylated 5hmC nucleobases to perform base excision or de-glucosylation, respectively. Brig3 generates abasic sites on non-glucosylated DNA, on phage DNA that was either first attacked by BapA or that naturally contains unmodified 5hmC nucleobases.

We incubated each resulting oligonucleotide overnight with either BapA or Brig3, purified the DNA products to remove protein and change the reaction buffer, and proceeded to a second 1-hour incubation with the other enzyme. As a control, we also carried out the second incubation with Brig1, which excises α-, but not β-, glucosyl-5hmC nucleobases (17). Samples were treated with NaOH and heat to promote β-elimination at abasic sites, and the cleaved oligonucleotides were separated by PAGE. Cleavage was observed when the modified oligonucleotides were first incubated with BapA and then Brig3, but not in the reverse order (Fig. 5B), and was more pronounced for α-than β-glucosylated substrates. We also observed an inhibitory activity of BapA on Brig1’s ability to generate abasic sites on the α-glucosyl-5hmC-containing oligonucleotide (Fig. 5B). These results suggest that BapA modifies glucosyl-5hmC nucleobases to generate Brig3’s substrate (5hmC) and at the same time eliminate Brig1’s substrate (α-glucosyl-5hmC). Therefore, we hypothesized that BapA removes α- and β-glucosyl groups from glucosylated 5hmC nucleobases present in DNA. To test this, we measured free glucose levels after incubation of the α- or β-glucosyl-5hmC-containing oligonucleotide, as well as a control containing an unmodified cytosine, with BapA. Glucose was quantified through the generation of NADH by glucose dehydrogenase, which in turn is detected via a quantitative luminescence assay (36). Compared to either lack of enzyme addition or treatment of the control oligonucleotide, we observed a significant increase in free glucose after incubation of the α-glucosylated substrates with BapA, and a small increase in the case of the β-glucosyl-5hmC oligonucleotide (Fig. 5C). This result demonstrates that BapA (i) releases glucose from glucosylated 5hmC nucleobases and (ii) is more active on α- than β-glucosylated 5hmC, consistent with Brig3 base excision following BapA activity (Fig. 5B). To test for the impact of this differential activity on anti-phage defense, we infected *E. coli* expressing both Brig3 and BapA with T4(Δ*a-gt*) and T4(Δ*b-gt*) phages and enumerated PFUs. We found that while the combined action of these enzymes completely prevented the detection of T4(Δ*b-gt*) plaques, T4(Δ*a-gt*) PFUs were decreased by two additional orders of magnitude when compared with propagation in cells producing Brig3 alone (three orders of magnitude when compared to a vector control), with a substantial level of phage propagation still taking place (Fig. S3C). However, when we over-expressed Brig3 and BapA using an arabinose-inducible promoter, we failed to detect T4(Δ*a-gt*) plaques (Fig. S3C). Together, these experiments are in line with in vitro results showing that BapA is less efficient at removing β-glucosyl groups from 5hmC.

We decided to confirm BapA activity using viral DNA, the actual substrate for the enzyme during infection. We purified DNA from different phages and measured glucose release after treatment with BapA. We found high levels of free glucose when the T4(wt) genome was used as substrate (Fig. 5D). In contrast, glucose release from the T4(5hmC) genome was similar to a control reaction that lacked DNA (Fig. 5D). To further investigate differential activity of BapA on α- and β-glucosyl groups, we performed the assay using genomic DNA from the T4(Δ*a-gt*) and T4(Δ*b-gt*) genomes, which display ∼75% α-glucosylation and ∼100% β-glucosylation of their 5hmC nucleobases, respectively. We detected high levels of glucose upon treatment with BapA of both genomes (Fig. S8E), with similar values for the release of α- and β-glucosyl groups that most likely reflect the saturation of the glucose signal in the assay. To test whether BapA can remove glucosyl groups from doubly glucosylated 5hmC sites (gentiobiosyl-5hmC) present in the T6 genome, we treated purified T6 phage DNA and tested for the release of glucose. We found low levels of glucose, similar to the levels in the control reactions in which either no DNA or T4(5hmC) phage DNA was used (Fig. 5D). We interpreted the lack of a significant signal as the inability of BapA to remove glucose from gentiobiosyl-5hmC, since only ∼3% of the 5hmC nucleobases are α-glucosylated, i.e., are suitable substrates (14). To corroborate this result, we performed the experiment using T6(Δ*g-gt*) DNA, harboring a deletion of the gene encoding for the second glucosyltransferase that leads to the generation of a genome that contains ∼75% of α-glucosyl-5hmC nucleobases. We detected high levels of free glucose upon incubation with BapA (Fig. 5D), data that is in line with our previous results using T4(Δ*b-gt*) DNA (Fig. S8E). Therefore, BapA cannot release glucose from more complex sugar modifications like gentiobiose.

We conclude that BapA is a glucosyl hydrolase that uses phage DNA containing α- and β-glucosyl-5hmC as substrates (with preference for the former isomer), generating 5hmC nucleobases in viral genomes for their excision by Brig3 (Fig. 5E). We wondered whether BapA could work independently of Brig3 and generate non-glycosylated phage genomes that could then be subject to 5hmC base excision. To test this, we propagated T4(wt), T4(Δ*a-gt*) and T4(Δ*b-gt*), as well as T4(5hmC) as a control, in an *E. coli* strain that harbors a plasmid for the isopropyl-β-D-1-thiogalactopyranoside-(IPTG-) inducible over-expression of BapA. The resulting phages were plaqued on a strain containing pBrig3 to enumerate PFUs (Fig. S8F). Compared to the plaquing of these phages of *E. coli* expressing the *brig3*-*bapA* operon (Figs. 1A and S3C), we found that initial passaging through cells expressing only BapA did not change the immunity provided by Brig3 alone, i.e., did not reduce PFUs for T4(wt) nor T4(Δ*a-gt*) phages (Fig. S8F).

Moreover, plaquing on bacteria expressing Brig1, which requires the presence of α-glucosyl-5hmC nucleobases to provide immunity, was not affected (Fig. S8G). These results suggest that BapA is less efficient than T4 glucosyltransferases in vivo and cannot generate phage genomes that contain a sufficiently low fraction of glucosylated 5hmC residues to enable Brig3 immunity or limit Brig1 defense. Instead, the data suggest that, to disrupt viral replication, Brig3 must generate abasic sites rapidly after BapA activity, before glucosyltransferases regenerate T4 glucosylation.

### An aspartic acid is required for BapA’s activity

We were interested in understanding the molecular mechanism of BapA activity but were unable to obtain its crystal structure. We therefore generated an AlphaFold 3 structural model of BapA bound to a 5hmC-containing dsDNA molecule (Figs. 6A and S9A; ipTM = 0.94, pTM = 0.94). In this model, BapA adopts a conformation with two central β-barrel domains surrounded by several α-helices. Positively charged surface patches form a continuous cleft that spans the linkers connecting each β-barrel to adjacent α-helices, with β-barrel 1 acting as the primary mediator of DNA interaction (Fig. S9B). This structure is characteristic of ASC-1 homology (ASCH) domain family proteins, known to bind nucleic acids and play essential roles in RNA metabolism (37). In particular, binding of DNA by the β-barrel 1 of BapA is similar to that of an ASCH protein from *Zymomonas mobilis* (ZmASCH, Fig. S9C; RMSD = 1.108 Å), which binds nucleic acids and promotes RNA cleavage through a structurally analogous cleft (38). The structural prediction indicates that the 5hmC nucleobase is flipped out of the DNA double helix and inserted into a putative catalytic pocket formed by the extended β1-α5 linker, where it is predicted to form a hydrogen bond between the hydroxyl group of the nucleobase and the carbonyl group of D101 side chain (Fig. 6B). The space left by the flipped base within the DNA is occupied by the Q252 side chain, which carbonyl group forms a hydrogen bond with the amino group of the complementary guanine base (Fig. 6B). If the model is accurate, mutations of these residues should affect BapA activity and therefore we tested the effect of D101N and Q252A substitutions in vitro and in vivo. We purified both mutant versions of the protein (Figs. S9D-E) and incubated them with oligonucleotide substrates harboring α-glucosyl-5hmC nucleobases to measure glucose release. We found that the D101N, but not the Q252A, mutation drastically affected BapA activity (Fig. 6C). We also tested Brig3 cleavage of the oligonucleotide substrate, after treatment with the mutant BapA proteins. We also detected a marked reduction of cleavage upon incubation with the D101N mutant, but the activity of the Q252A mutant was undistinguishable from that of wild-type BapA (Fig. 6D). Finally, we corroborated the importance of the D101 residue in vivo, after infecting *E. coli* cells expressing Brig3 and either wild-type or the D101N mutant BapA with T4(wt) (Fig. 6E). Together, our data demonstrates that BapA employs an aspartic acid, D101, to remove glucose residues from α- and β-glucosyl-5hmC nucleobases, most likely participating in the specific recognition of the modified base and/or the catalysis of glucose hydrolysis.

**Figure 6.**
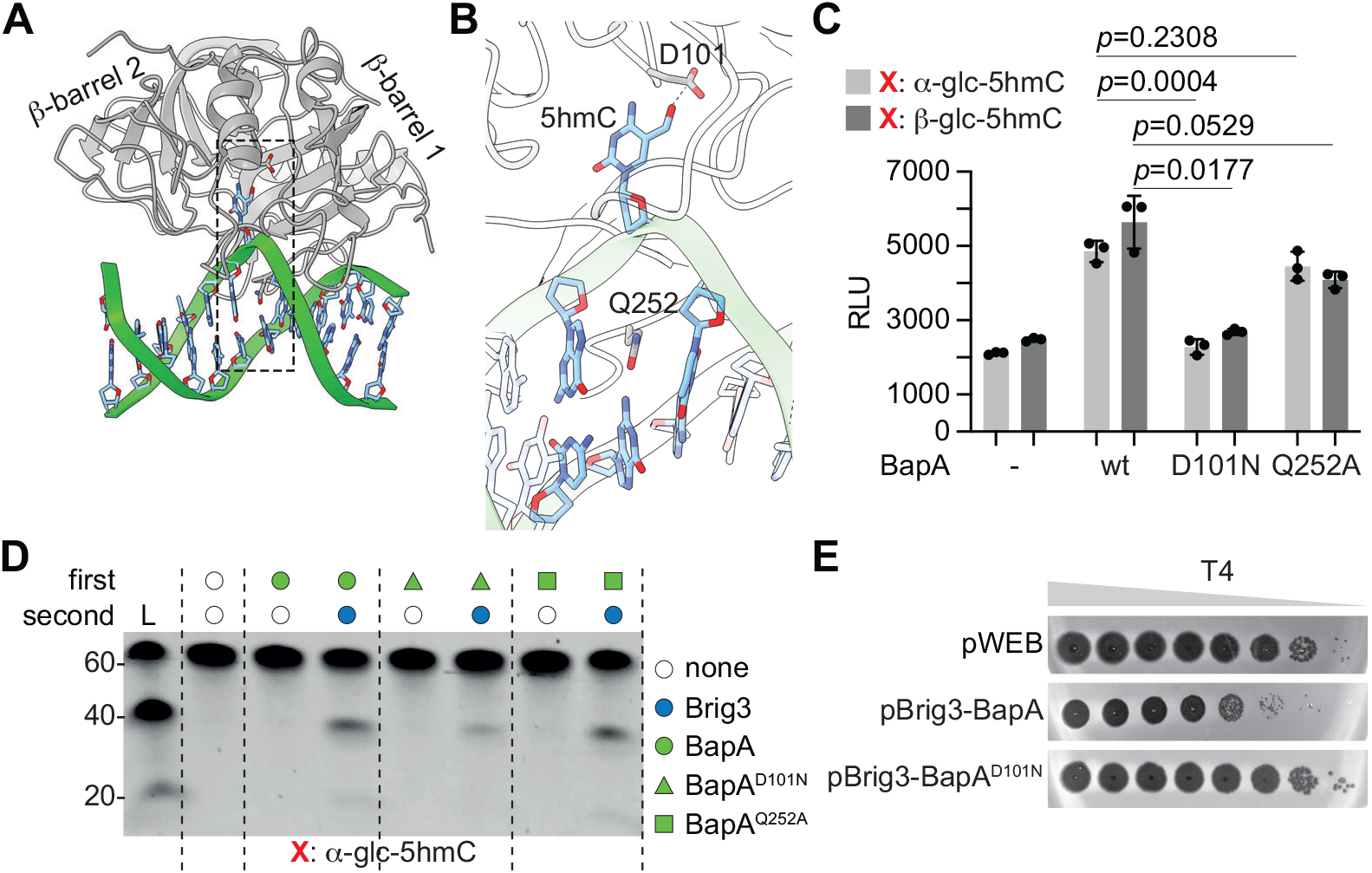
Aspartate 101 is essential for BapA activity. **(A)** AlphaFold 3 model of BapA with a dsDNA duplex harboring a 5hmC nucleobase, showing the two β-barrels present in the protein. The predicted aligned error (ipTM = 0.94, pTM = 0.94) supports a confident global fold. **(B)** Region highlighted by the inset in **(A)**. Residue D101 interacts with the 5hmC nucleobase. Residue Q252 is inserted between the base pairs flanking the flipped 5hmC nucleobase. **(C)** Luminescence assay for glucose detection after 37°C overnight incubation of oligonucleotides harboring α- or β-glucosyl-5hmC with either wild-type (wt) BapA, D101N or Q252A mutants, or no enzyme (−). RLU, relative luminescence units. Error bars represent the standard error of the mean; *p*-values are reported for a Ordinary One-Way ANOVA with multiple comparisons. **(D) PAGE of** α-glucosyl-5hmC oligonucleotides incubated at 37°C overnight, first with wild-type (wt) BapA, D101N or Q252A mutants, and then with Brig3. Samples were heated in the presence of NaOH for 30 minutes prior to electrophoresis. Gel was stained with ethidium bromide. L, ssDNA size ladder. **(E)** Ten-fold serial dilutions of T4 phage on lawns of *E. coli* EC100 that harbor the pWEB cosmid expressing Brig3 and wild-type or D101N BapA.

## DISCUSSION

In this work, we used an environmental DNA library to search for genes that prevent the propagation of a T4 mutant phage lacking glucosyltransferases and therefore harboring non-glucosylated 5hmC nucleobases in its genome. We found Brig3, which we characterized in vitro as a 5hmC-specific DNA glycosylase that generates abasic sites in the viral DNA (Fig. 5E). In vivo, Brig3 prevents the accumulation of phage DNA within infecting cells. Therefore, we propose a mechanism of immunity in which the introduction of multiple abasic sites throughout the viral genome (in dsDNA and/or ssDNA replication intermediates (39), since Brig3 can generate abasic sites on 5hmC-containing phage dsDNA and ssDNA oligonucleotides) interferes with phage replication and/or transcription and enables host survival. It is also possible that Brig3 leads to the spontaneous hydrolysis of the viral sugar-phosphate backbone due to the reactivity and instability of abasic sites (40). In addition, abasic sites lead to “transcriptional mutagenesis” that generates mutant phage transcripts due to the misincorporation of the base opposite to the abasic site in the template DNA strand, predominantly adenine misincorporation (41); such mutations could be detrimental to the viral lytic cycle. Brig3 provides defense against a wide range of T-even phages with partial glucosylation of their 5hmC nucleobases, but cannot interfere with wild-type T4, which lacks free 5hmC sites in its genome (7). Interestingly, *brig3* is part of a two-gene operon also containing *bapA*. In vitro, BapA is a glucosyl hydrolase that removes glucose groups from both α- and β-glucosyl-5hmC nucleobases (Fig. 5F). When co-expressed with Brig3 in vivo, BapA extends the defense range to include phages with fully glucosylated genomes, such as wild-type T4.

The *brig3*-*bapA* operon is located on an anti-phage defense island present within the same environmental DNA sample that contained the α-glucosyl-5hmC DNA glycosylase Brig1 (17) and therefore may be evolutionary related at two levels. First, since both systems possess residual activity on uracil, we believe that they emerged from uracil DNA glycosylases involved in DNA repair that were repurposed to excise non-canonical nucleobases within phage genomes to provide defense. Second, similarly to *E. coli* restriction endonucleases McrBC (13) and GmrSD (42), which recognize 5hmC and glucosyl-5hmC, respectively, Brig1 and Brig3 may have also evolved to recognize increasingly complex DNA modifications of the viral genome in the context of the bacteria vs phage arms race. It is plausible that, upon the introduction of 5hmC in the genome of T-even-related phages to limit restriction, bacteria responded with the development of Brig3 to regain immunity. As phages glucosylated 5hmC bases to further limit the access of DNA processing enzymes involved in bacterial defense to their genomes, hosts counteracted either with a new DNA glycosylase with specificity towards the new, hypermodified base, or with the incorporation of BapA to complement Brig3 activity. From our admittedly limited data, we conclude that this second strategy can provide more efficient immunity, both qualitatively and quantitatively. On one hand, the combination of Brig3 and BapA restricts infection by phages that contain 5hmC, α- or β-glucosyl-5hmC, as opposed to Brig1 that acts only on α-glucosylated viral genomes. In addition, when both systems are expressed using the same plasmid and arabinose-inducible promoter, Brig3-BapA lowers the PFU count of wild-type T4 to undetectable levels (Fig. 1C), whereas Brig1 reduces propagation of this phage by 5 orders of magnitude (17). The relatively high frequency of phages that escape Brig1 immunity is due to the presence of mutations that inactivate the *a-gt* gene in the viral population, which is not essential at least when using hosts related to the laboratory strain *E. coli* K-12 lacking the McrBC restriction system. In contrast, the only possibility to avoid base excision by the Brig3/BapA system requires the accumulation of inactivating mutations in *denA*/*denB*, *alc*, *gp56*, and *gp42* genes to prevent the incorporation of 5hmC into the T4 genome (43). This is highly unlikely and therefore explains the absence of “escaper” plaques in the presence of Brig3.

Our crystal structural data confirmed a base flipping mechanism that is widely conserved for the catalysis of base excision, with residue E55 directly recognizing the hydroxymethyl group of the flipped 5hmC base to provide the specificity of the reaction, and residue N225 replacing the excised base within the DNA double helix. This mechanism of base excision is similar to that of other DNA glycosylases involved in DNA repair in both prokaryotic and eukaryotic cells, where catalysis relies on a pre-organized active site that stabilizes a flipped base and positions the deoxyribose for glycosidic bond cleavage. Acidic residues activate a water molecule for nucleophilic attack, for example D64 in the *E. coli* uracil DNA glycosylase (UDG) (44), D145 in the human UDG homolog (45), or the K249-D268 pair in human 8-oxoguanine DNA glycosylase 1 (OGG1) (46). Base excision is generally coupled to the insertion of an amino acid side chain into the DNA duplex, such as the conserved L272 in human UDG (47), to maintain base stacking on the opposing strand and stabilize the distorted DNA geometry. Our findings therefore demonstrate that Brig3 preserves the core catalytic logic of DNA glycosylases while adapting substrate recognition to selectively target a modified viral nucleobase.

Despite several attempts, we were unable to obtain the crystal structure of Brig1 to compare it with that of Brig3. Therefore, we used AlphaFold 3 (48) to obtain a model for the α-glucosyl-5hmC DNA glycosylase bound to a DNA substrate containing a 5hmC base (ipTM = 0.91, pTM = 0.93), since it is currently not possible to model α-glucosyl-5hmC molecules in AlphaFold 3. We then compared the Brig1:DNA structural prediction to our Brig3^D127N^:DNA crystal structure. We found a close alignment for both structures (Fig. S10A; RMSD = 1.075 Å), with the DNA duplex is positioned in an almost identical cleft in both proteins and only minor shifts in the DNA backbone caused by conformational differences in the N-terminal helices. Examination of the catalytic pockets showed that, consistent with the need to accommodate and recognize a larger base, Brig1 contains a more open and expanded active site, formed by Q81, W122, and Q147 (Fig. S10B). In contrast, Brig3 relies on E55, W100, and N127 in its active site to coordinate 5hmC alongside Mg²⁺ ion and a catalytic water molecule (Fig. S10B). These observations suggest the existence of a shared mechanism of substrate engagement within the Brig family.

Our work expands the repertoire of DNA glycosylases involved in anti-phage defense, solidifying base excision as an efficient immune mechanism employed by bacteria against phages that carry modified genomes. It also highlights the advantages of the use of environmental DNA libraries to discover uncharacterized anti-phage defense systems. This method not only enables access to immunity genes currently hidden in the “microbial dark matter” (49) of uncultured, yet to be sequenced, bacteria living in diverse and remote environments, but also enables a targeted search for mechanisms of defense that attack specific aspects of phage biology, for example, by infecting the library with mutant phages such as T4(5hmC). More importantly, our findings uncovered an immune strategy in which base-modifying enzymes act on viral DNA to increase the range of phage genomes that are attacked by DNA glycosylases.

## METHODS

### Bacterial strains and growth conditions

Cultivation of *E. coli* strains used in this study were carried out in lysogeny broth (LB) at 37°C with shaking. Overnight cultures were inoculated from single bacterial colonies or glycerol stocks. LB media was supplemented with antibiotics where needed.

Chloramphenicol at 12.5 μg/mL (for cosmids) or 25 μg/mL (for plasmids), spectinomycin at 50 μg/mL, kanamycin at 50 μg/mL, ampicillin or carbenicillin at 100 μg/mL. Miniprepped plasmids (prepared by QIAprep Spin Miniprep Kit, QIAGEN, Cat# 27106) were cloned into electrocompetent *E. coli* EC100 cells (Lucigen, Cat# EC10010), *E. coli* BL21 (D3) (NEB, Cat# C2527H) or *E. coli* 5-alpha (Cat# C2987H). Transformations were performed by electroporation (1 mm Bio-Rad Gene Pulser cuvette at 1.8 kV). The list of strains used in this study are available in the Supplementary Data File.

### Plasmid construction

For plasmid construction details, refer to the Supplementary Data File.

### Gibson assembly

For standard Gibson assemblies (50), 25-100 ng of the largest dsDNA fragment was combined with equimolar volumes of the smaller fragment(s) in a total volume of 5 μL in nuclease-free water. Reaction mixtures were prepared on ice and mixed with 15 μL of Gibson assembly master mix, pipette mixed and incubated at 50°C for 1 hour in a thermal cycler. Gibson reactions were transformed into electrocompetent or chemically competent *E. coli* cells by following a standard transformation protocol.

For NEBuilder® HiFi DNA assembly we modified the standard protocol and used the following for each reaction: 1.5 μL NEBuilder® HiFi DNA Assembly master mix, 1.5 μL of purified DNA from the fragment to be cloned, 0.3 μL of purified DNA from the vector. The reactions were mix and incubated at 50°C for 30 mins. For transformation, 1 μL of the reaction was electroporated directly into *E. coli* EC100 cells (Lucigen, Cat# EC10010) and a standard transformation protocol for electrocompetent cells was followed.

### Preparation of phage stocks

T4 was a generous gift from Bruce Levin. T2 and T6 phages were purchased from ATCC. T4 and T6 mutants were generated in our previous published study (17). Phages and phage mutants are listed in the Supplementary Data File. Phages were first grown up in 10 mL cultures of exponentially growing EC100 cells at OD_600_∼0.3. The phage-infected cultures were incubated at 37°C with shaking overnight. Tubes were then spun down at 15,000 x *g* for 10 minutes at 4°C. Phage-containing supernatants were filtered using Acrodisc 13 mm SUPOR 0.45 μm syringe filters (Pall, 4604) into 15 mL conical tubes and stored at 4°C. Final phage stocks were titered on top agar lawns of *E. coli* EC100.

### Plaque assays and efficiency of plaquing analysis

Overnight cultures were started from single colonies in LB medium supplemented with appropriate antibiotics. Top agar lawns of *E. coli* were prepared by mixing 100 μL of overnight culture with 6 mL of LB top agar (LB broth Lennox base, 0.5% agar) supplemented with appropriate antibiotics. Top agar mixtures were poured over LB agar plates supplemented with appropriate antibiotics. For some experiments, 0.2% L-arabinose was included in the overnight media as well as in the LB top agar and the LB agar plate. Plates were dried at room temperature for 20 minutes for the top agar to solidify with an open flame. Serial dilutions of phage stock were prepared and 2.5 μL of each phage dilution was spotted on top agar using a multichannel pipette. After, plates were incubated at 37°C overnight after drying at room temperature until the plates were completely dry. Overnight plaque assays were imaged the next day (∼16-24 hours after infection) using the FluorChem HD2 system (ProteinSimple). Plaque assay images were all auto-contrasted using Adobe Photoshop to give clearer images. In some cases, image brightness was modified using Adobe Photoshop.

### Functional selection of a T4(5hmC)-resistant clone in the AZ52 soil DNA library in *E. coli*

The DNA library we used was generated using DNA extracted from an arid soil sample collected in Arizona (19). The library, AZ52, is comprised of large ∼40 kb DNA fragments from soil microorganisms cloned into a pWEB-TNC cosmid (abbreviated “pWEB” in the text and figures). The cosmids were transformed into *E. coli* EC100 cells (Lucigen), generating a soil DNA library with approximately 20 million clones, divided into megapools (17, 19, 51).

Each clone contains a cosmid with a soil DNA insert, which carries genes from unknown soil microorganisms. Genes from the soil samples can therefore be expressed heterologously in our library system. We performed our environmental eDNA functional screen to detect new bacterial immune defense systems (detailed in our previous study (17)), using phage T4 Δ*a-gt* Δ*b-gt*. To grow up libraries, we scraped frozen library stocks of *E. coli* EC100 carrying megapools 3-16 of the AZ52 DNA library into separate tubes with 10 mL LB supplemented with 12.5 μg/mL chloramphenicol and grew cultures overnight at 37°C with shaking. The next day, we infected *E. coli* EC100 overnight cultures with T4Δ*a-gt* Δ*b-gt* at a multiplicity of infection (MOI) of 10. Infections were performed in 6 mL LB top agar with 500 μL of overnight culture mixed with phage at MOI 10 on LB agar plates, supplemented with 12.5 μg/mL chloramphenicol. We incubated plates at 37°C for 72 hours and then inspected surviving colonies within top agar infections. We found that megapools 3, 9 and 10 showed an increased number of surviving colonies upon T4(5hmC) infection compared to an infection of *E. coli* EC100 cells carrying an empty pWEB-TNC cosmid (control).

To eliminate false positive clones, we extracted pooled cosmid DNA from the surviving colonies on the enriched plate. To do this, we scraped top agar with surviving colonies into a 50 mL conical tube, melted the top agar in a 98°C heating block for 10-15 minutes until the top agar was completely melted, and then centrifuged the tube at ∼4000 x *g* for 5 minutes at room temperature to collect a cell pellet from which surviving cosmids were isolated using the QIAprep Spin Miniprep Kit (QIAGEN, Cat# 27106). The miniprepped cosmid pool was then transformed into 50 μL of electrocompetent *E. coli* EC100 cells (Lucigen) through electroporation (1 mm Bio-Rad Gene Pulser cuvette at 1.8 kV) and recovered in 1 mL SOC medium. After 1.5 hours of recovery, cells were assayed for transformation efficiency by pipetting ten-fold serial dilutions of the transformation culture on to an LB agar plate supplemented with 12.5 μg/mL chloramphenicol. While the plate was grown overnight at 37°C, the remaining transformation culture was stored overnight at 4°C. The next day, based on the calculated transformation efficiency, the transformation culture was spread onto ten 15 cm LB agar plates supplemented with 12.5 μg/mL chloramphenicol, plating for ∼30,000 colonies on each plate, for a total of ∼300,000 colonies. Plates were incubated overnight at 37°C and the next day colonies from all ten plates were scraped into 15 mL LB, vortexed and inverted to mix, and then diluted to OD_600_ = 20. The OD_600_ = 20 colony mixture was then mixed 1:1 with 50% glycerol to make a −80°C freezer stock of a 1X phage-enriched DNA library for AZ52 megapools 3,9 and 10.

We did not enrich this library further by re-infection. We sampled colonies from the 1x-enriched library for anti-phage immunity by streaking the library to single colonies on an LB agar plate supplemented with 12.5 μg/mL chloramphenicol. Multiple colonies were grown overnight in LB supplemented with 12.5 μg/mL chloramphenicol at 37°C with shaking. Colonies were assayed for anti-phage immunity using plaque assays (described above) with T4 Δ*a-gt* Δ*b-gt*. Cosmids were isolated from the T4(5hmC)-resistant clones using the QIAprep Spin Miniprep Kit (QIAGEN, Cat# 27106) and sent for “huge plasmid sequencing” by Plasmidsaurus. Sequencing revealed 3 resistant colonies contained a different metagenomic DNA insert. These colonies were frozen at −80°C (900 μL culture + 100 μL DMSO) for use in future experiments.

### Cosmid sequencing, assembly, and gene annotation

Cosmid DNA was extracted using the QIAprep Spin Miniprep Kit (QIAGEN, Cat# 27106). DNA was sequenced by Plasmidsaurus. The sequence of the cosmid harboring Brig3 was deposited on GenBank, accession number PX942695. PADLOC (52) (https://padloc.otago.ac.nz/padloc/) was used to predict ORFs within the metagenomic DNA insert of the assembled cosmid genome. Predicted ORFs were then run through NCBI PSI-BLAST (https://blast.ncbi.nlm.nih.gov/Blast.cgi?PAGE=Proteins), HHpred (22) (https://toolkit.tuebingen.mpg.de/tools/hhpred) and Alphafold 2 (23, 53) https://colab.research.google.com/github/sokrypton/ColabFold/blob/main/AlphaFold2.ipynb) and DALI (24) (http://ekhidna2.biocenter.helsinki.fi/dali/) to predict protein function where possible. These predictions helped to design the fragments for subcloning of the cosmid and identifying the anti-T4(5hmC) phage defense system.

### Subcloning of the T4(5hmC)-resistant cosmid to identify the phage system

We subcloned 6 DNA fragments (1–6) from the entire length of the metagenomic insert sequence in the resistant clone (Fig. S1A). DNA fragments were amplified using 10 ng of cosmid DNA as template for PCR amplification using Phusion High-Fidelity DNA Polymerase (Thermo Scientific, Cat# F530L) with 1 M betaine (Sigma-Aldrich, Cat# B0300) and 1 μL DMSO in a 50 μL PCR reaction. Fragments were cloned into PCR-amplified pWEB-TNC cosmid backbones using NEBuilder® HiFi DNA Assembly Master Mix (NEB, Cat# E2621L). Reactions for assembly with NEBuilder® HiFi were carried out as described above. Transformations were performed by electroporation (1 mm Bio-Rad Gene Pulser cuvette at 1.8 kV). Cells were then plated on LB agar supplemented with 12.5 μg/mL chloramphenicol and incubated overnight at 37°C. The next day 4 colonies were picked, grown overnight in LB supplemented with 12.5 μg/mL chloramphenicol and their cosmids miniprepped using the QIAprep Spin Miniprep Kit (QIAGEN, Cat# 27106). Miniprepped cosmids were sent to Plasmidsaurus. Colonies that harbored cosmids with correct insert fragments were then assayed for immunity against phage T4(5hmC) using plaque assays (see above). Plaque assays identified Fragment 6 as the fragment harboring immunity. Fragment 6 was further subdivided into Fragments 6-1, 6-2, 6-3 and 6-4, cloned and tested for immunity as described above. Fragment 6-1, containing Brig3, was identified as the minimal DNA fragment carrying anti-T4(5hmC) immunity.

Brig3 protein structure was predicted at this early stage by AlphaFold 2 (23, 53) https://colab.research.google.com/github/sokrypton/ColabFold/blob/main/AlphaFold2.ipynb).

### Genomic integration of Brig3 and BapA

λ Red recombineering (54) was used to generate the *E. coli* EC100 strain gBrig3 and, which contains *brig3* and its putative promoter region integrated into the *attnTn7* site(26). An overnight culture of *E. coli* EC100 carrying the pAM38(*red*) plasmid with chloramphenicol resistance was diluted and grown to OD_600_ ∼ 0.3 and then induced with 0.2% L-arabinose until OD_600_ ∼ 1–1.2. Cells were made electrocompetent by washing twice with cold water and electroporated (1 mm Bio-Rad Gene Pulser cuvette at 1.8 kV) with a DNA construct carrying either the Brig3 gene or the Brig3-BapA genes, and the Kanamacyn resistance gene, with ∼50 bp homology upstream and downstream of the attTn7 site. After ∼2 h of recovery, cells were plated on LB agar plates with kanamycin at 50 μg ml^−1^ to select for genomic integration. Gene integration was confirmed by PCR. After confirmation, strains were grown overnight in LB with kanamycin at 50 μg ml^−1^ (but no chloramphenicol which selects for the plasmid) and with 0.2% L-arabinose induction. Without antibiotic selection, induced plasmid was lost due to toxicity from λ Red overexpression. Strains were frozen at −80 °C (900 μl culture + 100 μl DMSO) and struck out on appropriate antibiotic plates to confirm both genomic integration of Brig3 or Brig3-BapA and loss of the recombineering plasmid.

### Western Blot of His-Tagged Brig3 Protein Expression in *E. coli*

*E. coli* EC100 cultures (100mL each) expressing Brig3-His_6_ under its native promoter either in a plasmid (pBrig3-His_6_) or in the genome (gBrig3-His_6_), or a control plasmid (pWEB-TNC) were grown in LB medium, at 37°C with shaking overnight. Cells were harvested by centrifugation 4,500 rpm x 10 min at 4°C. Cell pellets were resuspended in 20 mL of lysis buffer (300mM Tris, 200mM NaCl) and lysed by sonication (50% amplitude, 10s pulses, 20s breaks). Lysates were mixed with an equal volume of 2× Laemmli sample buffer containing β-mercaptoethanol and denatured by heating at 98°C for 5 min. Purified Brig3-His_6_ protein was serially diluted and used as a positive control, mixed 1:1 with 2× Laemmli sample buffer and denatured as above. The final loading concentration was 4.5 ng per lane. Samples were loaded on a 4-20% polyacrylamide gel (BioRad) alongside a molecular weight marker (Pierce Prestained Protein MW Marker, Thermo Fischer) for SDS-PAGE and transferred to a nitrocellulose membrane (0.2 μm, BioRad) using a Trans-Blot Turbo transfer system (BioRad). After blocking the membrane in 5% milk in PBST (PBS, 0.1% Tween20 (Sigma-Aldrich)) for 1h, His_6_-Brig3 was probed with primary mouse THE His Tag anti-His_6_ antibody (GeneScript, AA00186) 1:5000 in PBST with 3% BSA, overnight at 4°C. The membrane was incubated with the secondary goat anti-mouse IgG (H+L) HRP antibody (ThermoFischer, 31430) diluted 1:10000 in 5% milk in PBST for 1 hour. HRP signal was detected with Clarity Western ECL substrate (BioRad) and ImageQuant 800 (Amersham).

### Phage DNA extraction

Phage genomic DNA was performed extracted from phage capsids using a previously described protocol (55). In brief, three tubes (or more depending on the phage titer) of 450 μL of a phage stock were first treated with DNase I (Invitrogen, Cat# 18068015) and RNase A (Promega, Cat# A7973) in DNase I buffer (20 mM Tris-HCl, pH 8, 2 mM MgCl_2_), the reaction stopped with EDTA (Invitrogen, Cat# AM9260G), then capsids digested with Proteinase K (NEB, Cat# P8107S), and finally phage genomic DNA extracted using the DNeasy Blood & Tissue kit (QIAGEN, Cat# 69504). DNA was quantified using the Qubit™ dsDNA HS Assay Kit and assessed for quality using a nanodrop spectrophotometer.

### Purification of Brig3 for *in vitro* assays

The *brig3* gene was cloned into pET21a and added a His_6_ tag immediately after the native C-terminal of Brig3 (pET21a-Brig3-6xHis). The insert was verified by DNA sequencing. *E. coli* strain BL21 (DE3) was used for protein expression. Cells were grown in LB medium with 100 μg/mL ampicillin at 18°C. 0.5 mM IPTG was added to induce protein expression when OD_600_∼0.7, followed by further growth at 37°C for 2 hours. Cell pellets were resuspended in Ni column buffer A (50mM HEPES, 1 M NaCl, 5% glycerol, 1 mM TCEP, pH 7.5) with complete mini protease inhibitor cocktail (Roche), one tablet/1 L culture. After adding lysozyme to a final concentration of 200 mg/mL, the mixture was sonicated 3 times for 1 minute each, then centrifuged at 20,000 rpm in an SS-34 rotor for 1 hour. The supernatant was filtered and loaded onto a Ni gravity column (ThermoScientific, HisPur™ Ni-NTA Resin, Cat#88222), and eluted with a gradient of 0 to 100% buffer B (Ni buffer A plus 300 mM imidazole, pH 7.5). The Brig3 containing fraction was further purified using Size Exclusion Chromatography with Buffer A.

### qPCR of phage DNA replication

Overnight cultures of *E. coli* EC100 cells carrying pWEB-TNC or pBrig3 were diluted 1:50 in 10 mL of LB medium supplemented with 12.5 μg/mL chloramphenicol. After 1 hour 45 minutes of growth, OD_600_ was measured, and the culture was normalized to OD_600_ = 0.3. 700 μL of culture was dispensed between multiple 1.5 mL Eppendorf tubes, corresponding to three replicates and multiple timepoints. These 700 μL cultures were infected with phage T4(5hmC) at MOI 1 and incubated at 37°C with shaking for specified timepoints. At each timepoint (2,4,8 minutes post infection), samples were removed from the incubator and tubes spun down at 15,000 rpm for 1 minute in a tabletop microcentrifuge at 4°C. Supernatants were removed and cell pellets immediately frozen down at −80°C for DNA extraction.

Total DNA was extracted from frozen *E. coli* cell pellets using the Promega Wizard® Genomic DNA Purification Kit (Promega, Cat# A1125) following the protocol for Gram-negative bacteria. Extracted DNA was quantified using the Qubit™ dsDNA HS Assay Kit and each sample was normalized to 4 ng/μL. A total of 32 ng DNA was used as input for qPCR, performed using Fast SYBR Green Master Mix (Applied Biosystems, Cat# 4385612) and the QuantStudio® 3 Real-Time PCR System (Applied Biosystems) with primer pairs AA870/AA871 (T4 *gp43* target), AA872/AA873 (T4 *gp34* target) and AA387/AA388 (*E. coli* K-12 MG1655 *dxs* control) used also in a previous study (17). For qPCR data analysis, ΔΔCt values were calculated for the two T4(5hmC) qPCR targets for each replicate at each timepoint. Fold-change values were then calculated for each replicate relative to the mean ΔΔCt value for cells carrying pWEB-TNC infected with T4(5hmC) phage at the earliest timepoint post infection for a each experiment. The mean fold change of three biological replicates was plotted for each timepoint post infection.

### Generation of glucosylated ssDNA

ssDNA oligonucleotides were alpha- or beta-glucosylated for *in vitro* assays with BapA following the protocol reported previously (17). hmdC_18 and hmdC_60_MfeI, 18 mer and 60 mer oligonucleotides, which contain one single 5hmC residue (Supplementary Data File) were glucosylated using either T4 beta-glucosyltransferase (b-GT) from NEB (Cat # M0357S) or alpha-glucosyltransferase (a-GT) purified previously (17). 100 μM of Substrate ssDNA was mixed at a 1:1 molar ratio with a-GT or b-GT in 1X NEBuffer 4 (50 mM potassium acetate, 20 mM Tris-acetate, 10 mM magnesium acetate, 1 mM DTT, pH 7.9) supplemented with 2 mM UDP-Glucose (NEB, supplied with NEB T4 beta-glucosyltransferase (b-GT)). All samples were incubated at 37°C overnight, then purified with an oligonucleotide cleanup kit (Zymo Research, Oligo Clean & Concentrator Kit, Cat# D4061) according to the manufacturer’s instructions. Final glucosylated oligonucleotide concentration was measured using the Qubit™ dsDNA HS Assay Kit and assessed for quality using a nanodrop spectrophotometer for use in following experiments.

### Annealing of ssDNA oligonucleotides

To generate dsDNA substrates for DNA glycosylase assays, complementary ssDNA oligonucleotides were annealed. In brief, 1:1 molar ratios of top and bottom strand complementary ssDNA oligonucleotides (25 μM each) were mixed in a 60 μl reaction containing NaCl to a final concentration of 100 mM. The reaction was heated at 80 °C for 20 min in a thermal cycler and then allowed to cool very slowly to room temperature. Annealed oligonucleotides were purified using an oligonucleotide cleanup kit (Zymo Research, Oligo Clean & Concentrator Kit, D4061) according to the manufacturer’s instructions.

### DNA glycosylase assays with ssDNA or dsDNA oligonucleotides: detection by NaOH-or endonuclease IV-mediated cleavage of the abasic sites

DNA glycosylase reactions were carried out in a reaction buffer containing 45 mM HEPES, pH 7.5, 0.4 mM EDTA, 2% glycerol, 1 mM DTT and 50 mM KCl, in a total reaction volume of 50 μL. The final ssDNA and dsDNA concentrations were 1 μM in every experiment (oligonucleotides used in this study can be found in the Supplementary Data File). Brig3 or Brig3 active site mutants (D127A, D127N or Y97) were added at different final concentrations: 1 μM (Fig 3B and S4G), 100nM (Fig. S4D, and Fig. S7F), or increasing concentrations for Brig3 titration (Fig. 3C and S4C). As a control, 2 μL (10 units) of hSMUG1 (NEB, Cat# M0336S) was added to some samples in Fig. 3B. Reactions were incubated at 37°C for 30 minutes. A second matched set of samples was treated with NaOH after initial incubation, 25 μL of 0.5 M NaOH was added to each 50 μL sample and then heated at 90°C for 30 minutes. For Fig S7F and S4H, oligonucleotide cleanup was also performed after NaOH and heat treatment to precisely visualize band cleavage and the size of the fragments generated (Zymo Research, Oligo Clean & Concentrator Kit, Cat# D4061) according to the manufacturer’s instructions. Samples were eluted from the cleanup columns in 15 μL nuclease-free water.

For all gels, 5 μL of each sample was mixed with loading dye and loaded onto a 10% TBE gel (Invitrogen, Cat# EC62755BOX) and electrophoresed at 140 V for 35 minutes. Gels were stained with 2 μg/mL ethidium bromide for 20 minutes, extensively rinsed with distilled water (3X for 10 minutes each), and then scanned using the Amersham ImageQuant 800 set to UV fluorescence. For gels in Fig 3B, 3C, S4C, the TriDye™Ultra Low Range DNA Ladder (NEB, Cat# N0558S) was used. For gels in Fig S4D and S7F DNA ladders were made by mixing 20-, 40- and 60-bp ssDNA oligonucleotides (Supplementary Data File) and loading them onto their corresponding gels at ∼100 ng each oligonucleotide per load.

For abasic site detection by NEB endonuclease IV (Endo IV), DNA glycosylase reactions were set up as described above and incubated with Brig3 for 30 minutes. After incubation, one sample was treated with NaOH as described above and purified using the Zymo Research Oligo Clean & Concentrator Kit (Cat# D4061) according to the manufacturer’s instructions. Another sample was processed directly using the oligonucleotide cleanup kit, then the purified sample was incubated at 37°C for 4 hours in a 50 μL reaction with 1X NEBuffer 3 (100 mM NaCl, 50 mM Tris-HCl, 10 mM MgCl_2_, 1 mM DTT, pH 7.9), with 50 units of NEB Endo IV (5 μL; 10 units/μL; Cat# M0304S). After 4 hours, the reaction was purified using the Zymo Research Oligo Clean & Concentrator Kit according to the manufacturer’s instructions. The control sample with no enzyme and just oligonucleotide was cleaned up after the initial overnight incubation. The second control sample with oligonucleotide and no enzyme also underwent NaOH treatment as mentioned above and later cleaned up. All the purified samples were then loaded onto a 10% TBE gel (Invitrogen, Cat# EC62755BOX), electrophoresed at 140 V for 35 minutes, stained with ethidium bromide as described above and imaged with the Amersham ImageQuant 800 set to UV fluorescence.

### Reversed-phase liquid chromatography and high resolution/high mass accuracy mass spectrometry of hSMUG1- and Brig1-treated ssDNA oligonucleotides

DNA glycosylase reactions were carried out in a reaction buffer containing 45 mM HEPES, pH 7.5, 0.4 mM EDTA, 2% glycerol, 1 mM DTT and 50 mM KCl, in a total reaction volume of 50 μL. Reactions were performed with 18mer ssDNA oligonucleotides: dC_18 and hmdC_18 (Supplementary Data File). The final ssDNA concentration in each reaction was 2 μM. Brig3 was added to a final concentration of 1μM. A no-enzyme reaction was used as a negative control. 2 x 50 μL reactions were set up for each condition. Reactions were incubated overnight at 37°C. After overnight incubation, samples were processed with an oligonucleotide cleanup kit (Zymo Research, Oligo Clean & Concentrator Kit, Cat# D4061) according to the manufacturer’s instructions.

For LCMS analysis, 3 μL (corresponding roughly to 18 pmol) was injected and separated on a 2.1mm*150mm, 5µm C18 column (Acclaim 120, Thermo Scientific) at 150µL/min. Solvent A was 100mM Hexafluoroisopropan-2-ol (Sigma), 15mM triethylamine (Sigma) in water, and solvent B was 100% methanol (Optima, Fisher Scientific). Oligos were resolved across a 15 minute gradient, increasing from 10% B to 50% B over 15 minutes. The gradient was delivered using a Dionex3000 NCS3500RS loading pump. Data acquisition was done using a Q-Exactive HF mass spectrometer (Thermo) operating in negative-ion mode, scanning from 500-1400 m/z at a resolution of 120000.

Raw data was converted to peaklists by selecting a representative swath of the chromatogram and exporting the summed spectra to text files. The text files were processed by UniDec GUI v.1.0.10 deconvolution software (56) in batch format using settings for low-resolution data.

### DNA glycosylase assays with phage and cosmid DNA

All reactions were performed in 50 μL reaction volumes in a reaction buffer containing 45 mM HEPES, pH 7.5, 0.4 mM EDTA, 2% glycerol, 1 mM DTT and 50 mM KCl.

Assays were performed by incubating 12.5 ng of extracted phage genomic DNA from capsids or miniprepped pWEB-TNC cosmid DNA with varying concentrations (2-800 nM) of purified Brig3. Reactions were incubated in a thermal cycler at 37°C for 30 minutes. Reactions were then mixed with 10 μL of purple 6X loading dye with no SDS (NEB, Cat #B7025S) and the entire reaction volume was loaded onto a 1% agarose gel containing ethidium bromide. The gel was run for 70 mins at 85 V at room temperature and then imaged using a UV gel imager (Amersham ImageQuant 800 set to UV fluorescence). The DNA ladder used was 1 kb Plus DNA ladder from NEB (NEB, Cat #N3200S).

### Two-step incubation assays with Brig3 and BapA

Alpha- and beta-glucosylated hmdC_60_MfeI 60-mer oligonucleotides were used in these experiments as indicated. All reactions were carried out in a reaction buffer containing 45 mM HEPES, pH 7.5, 0.4 mM EDTA, 2% glycerol, 1 mM DTT and 50 mM KCl, in a total reaction volume of 50 μL. For the first incubation, 1uM of oligonucleotides were incubated with either 100nM of BapA (or BapA mutants in Fig. 6D), 100nM of Brig3, 1μM of Brig1 or no enzyme as a control. Incubations were carried out at 37°C overnight and samples were processed with an oligonucleotide cleanup kit (Zymo Research, Oligo Clean & Concentrator Kit, Cat# D4061) according to the manufacturer’s instructions, eluted in 10μl nuclease-free water. The purified oligonucleotide product was used for the second incubation. Reactions were set up with the same buffer described above using 10 μl of the oligonucleotide product from the first incubation and with either 100nM of BapA, 100nM of Brig3, 1uM of Brig1 or no enzyme as a control. Reactions were incubated at 37°C for 1 hour and then 25 μL of 0.5 M NaOH was added to each 50 μL sample and then heated at 90°C for 30 minutes.

Samples were processed with an oligonucleotide cleanup kit (Zymo Research, Oligo Clean & Concentrator Kit, Cat# D4061) according to the manufacturer’s instructions after the second incubation. All the purified samples were then loaded onto a 10% TBE gel (Invitrogen, Cat# EC62755BOX), electrophoresed at 140 V for 35 minutes, stained with ethidium bromide as described for the glycosylase assays above and imaged with the Amersham ImageQuant 800 set to UV fluorescence. DNA ladders were made by mixing 20-, 40- and 60-bp ssDNA oligonucleotides (Supplementary Data File) and loading them onto their corresponding gels at ∼100 ng each oligonucleotide per load.

### Glucose detection assays with BapA and ssDNA oligonucleotides or phage DNA

Alpha- and beta-glucosylated 5hmC_18 18-mer oligonucleotides or extracted phage genomic DNA from capsids were used in these experiments as indicated. All reactions were carried out in a reaction buffer containing 45 mM HEPES, pH 7.5, 0.4 mM EDTA, 2% glycerol, 1 mM DTT and 50 mM KCl, in a total reaction volume of 50 μL. Oligonucleotides were used at 1μM final concentration and phage DNA at 100 ng per reaction. BapA was added to a final concentration of 100 μM per reaction and incubated at 37°C overnight. The next day, the whole 50 μL of the reaction was used for glucose detection using the Glucose-Glo™ Assay (Promega, Cat#J6021). The reactions were set up as directed in the protocol in a Nunc™ 96-well Optical-Bottom Black Microplate plate (ThermoFisher Scientific, Cat#165305). In brief, components were thawed and the following was added for each reaction: 50 μL luciferin detection solution, 0.25 μL reductase, 0.25 μL reductase substrate, 2 μL glucose dehydrogenase and 0.25 μL NAD. The plate was incubated at room temperature for an hour. Luminescence in RLU (relative fluorescence units) was obtained with a plate reader (Agilent, Biotek Synergy H1) as a single data point.

### BapA expression and purification

The *BapA* gene was codon-optimized and cloned into the pET21a vector with an N-terminal His-SUMO tag under ampicillin selection. The BapA mutants (D101N and Q252A) were generated using a high-fidelity PCR-based mutagenesis strategy, with the BapA–His plasmid serving as the template. Mutations were introduced with complementary primers carrying the desired nucleotide substitutions and amplified using PrimeSTAR Max DNA Polymerase (Takara R047B) according to the manufacturer’s instructions. Following amplification, the PCR reactions were treated with DpnI (New England Biolabs, R1076S) for 3 hours at 37 °C to selectively digest the methylated and hemimethylated parental plasmid DNA derived from E. coli. Newly synthesized PCR products, which lack adenine methylation, remain undigested and therefore enriched. The DpnI-treated reactions were directly transformed into E. coli DH5α competent cells, and colonies were screened by Sanger sequencing.

The sequence-confirmed plasmids were then transformed into BL21 (DE3) cells for recombinant expression in vitro by culturing in Terrific Broth (TB) medium and inducing with 0.5 mM isopropyl β-D-1-thiogalactopyranoside (IPTG) at 16 °C for 18 hours.

Purification of the BapA mutants followed the same procedures used for Brig2, while purification of tag-less BapA included one additional step: Ulp1 protease was added after Ni-NTA purification and incubated overnight at 4 °C, followed by passage through Ni-NTA resin to remove the His–SUMO tag and His-Ulp1 enzyme. The protein was then concentrated and run on a Superdex 75 Increase 10/300 GL size-exclusion column (GE Healthcare).

### AF3 analysis of BapA with 5hmc-containing DNA

The AF3-predicted model was analyzed to identify the BapA catalytic pocket and essential residues. Briefly, the prediction was performed online by inputting the full-length BapA sequence along with a 12-mer dsDNA containing hmC (5’ CGAGT/hmdC/GATGCG 3’) and its complementary strand (5’ CGCATCGACTCG 3’). The prediction was highly reliable, with most residues exhibiting pLDDT scores above 90, and high ipTM and pTM values of 0.94 each. The top five ranked models were very similar, and one was selected as a representative for catalytic pocket analysis.

### DNA solid-phase synthesis

DNA synthesis was performed on an ABI 392 nucleic acid synthesizer on a 1.0 µmol scale using phosphoramidite chemistry. The UltraMild canonical nucleoside building blocks of dA(pac), dC(pac), dG(dmf) and T, as well as CPG solid support (dG(iPr-pac) 3′-lcaa CPG 1000 Å), were purchased from ChemGenes. The modified phosphoramidite of 5-hydroxymethyl-dC(N,O-carbamoyl) was obtained from Glen Research. Detritylation, coupling, capping and oxidation reagents were dichloroacetic acid/1,2-dichloroethane (4/96), phosphoramidite/acetonitrile (100 mM) and benzylthiotetrazole/acetonitrile (300 mM), Cap A/Cap B (1/1) (Cap A: phenoxyacetic anhydride/tetrahydrofuran (200 mM), Cap B: *N*-methylimidazole (200 mM) and *sym*-collidine (200 mM) in tetrahydrofuran) and iodine (20 mM) in tetrahydrofuran/pyridine/H_2_O (35/10/5), respectively. Solutions of phosphoramidites and tetrazole were dried over activated molecular sieves (3 Å) overnight.

### Deprotection, purification and quantification of DNAs

For deprotection of unmodified DNAs the solid support was mixed with an aqueous ammonia solution (28%, 1.0 mL) for 4 h at room temperature. The supernatant was removed and the solid support was washed three times with H_2_O/ethanol (1.0 mL; 1/1 v/v); combined supernatant and washings were concentrated to a volume of *ca.* 1 mL.

Oligonucleotides modified with 5-hydroxymethyl-dC were treated with a fresh solution of 0.4 M NaOH in MeOH/H_2_O (1.0 mL; 4/1 v/v) and shaken at room temperature for 48 h. The supernatant was collected and neutralized with triethylammonium acetate/H_2_O (1.0 M, 1.0 mL, pH 7.4). The sample was desalted with size-exclusion column chromatography (GE Healthcare, HiPrep™ 26/10 Desalting; Sephadex G25) eluting with H_2_O; collected fractions were concentrated to a volume of *ca.* 1 mL.

The crude DNA was purified by anion exchange chromatography (Thermo Scientific Ultimate 3000 HPLC System) on a semipreparative Dionex DNAPac® PA-100 column (9 mm × 250 mm) at 80 °C with a flow rate of 2 mL/min (eluent A: 20 mM NaClO_4_ and 25 mM Tris·HCl (pH 8.0) in 20% aqueous acetonitrile; eluent B: 0.6 M NaClO_4_ and 25 mM Tris·HCl (pH 8.0) in 20% aqueous acetonitrile). Fractions containing DNA were evaporated and the residue redissolved in 0.1 M triethylammonium bicarbonate solution (10 to 20 mL), loaded on a C18 SepPak Plus^®^ cartridge (Waters/Millipore), washed with H_2_O, and then eluted with acetonitrile/H_2_O (1/1).

Crude and purified DNAs were analyzed by anion exchange chromatography (Thermo Scientific Ultimate 3000 HPLC System) on a Dionex DNAPac^®^ PA-100 column (4 mm × 250 mm) at 80 °C with a flow rate of 1 mL/min. A gradient of 0-40% B in 30 min was applied; eluent A: 20 mM NaClO_4_ and 25 mM Tris·HCl (pH 8.0) in 20% aqueous acetonitrile; eluent B: 0.6 M NaClO_4_ and 25 mM Tris·HCl (pH 8.0) in 20% aqueous acetonitrile. HPLC traces were recorded at UV absorption by 260 nm (Fig. S11A). DNA quantification was performed on an Implen P300 Nanophotometer.

### Mass spectrometry of oligonucleotides

DNA samples (*ca.* 200 pmol) were diluted with an aqueous solution of ethylenediaminetetraacetic acid disodium salt dihydrate (Na_2_H_2_EDTA) (40 mM, 15 µL). Water was added to obtain a total volume of 30 µL. The sample was injected onto a C18 XBridge column (2.5 µm, 2.1 mm × 50 mm) at a flow rate of 0.1 mL/min and eluted using gradient 0 to 100% B at 30 °C (eluent A: 8.6 mM triethylamine, 100 mM 1,1,3,3,3-hexafluoroisopropanol in H_2_O; eluent B: methanol). DNA was detected by a Finnigan LCQ Advantage Max electrospray ionization mass spectrometer with 4.0 kV spray voltage in negative mode (Fig. S11B).

### Protein crystallization

Apo-Brig3 crystals were grown at 16°C using the hanging-drop vapor diffusion method. In each drop, 1 µL of protein solution was mixed with 1 µL of reservoir solution, which contained 0.1 M citric acid (pH 5.0), 1 M lithium chloride, and 10% PEG6000. For the Brig3-5hmC-dsDNA complex, a 12-mer 5hmC-containing strand (5’CGAGThmCGATGCG) was incubated with its complementary strand (5’CGCATCGACTCG) in a 1:1 ratio. The DNA duplex was formed by self-annealing in an annealing buffer (25 mM HEPES, pH 7.5, 150 mM NaCl, and 2 mM MgCl_2_). The 5hmC-dsDNA was then incubated with wild-type Brig3 protein at a molar ratio of 1.3:1 at 4°C for 1 h, yielding a final protein concentration of 0.35 mM. Crystallization screening and optimization were performed following the same procedure used for apo-Brig3, yielding crystals in a solution of 0.1 M sodium citrate (pH 5.5) and 20% PEG3000. For the Brig3 D127N mutant complexed with 5hmC-dsDNA, a 13-mer 5hmC-containing strand (5’GCGAGThmCGATGCG) was used, along with the same complementary strand (5’CGCATCGACTCG). The strands were annealed using the same method as the wild-type complex, and the crystallization process was identical to that used for the apo-Brig3 and Brig3-5hmC-dsDNA complexes, resulting in crystals under the same conditions.

### Data collection, structure determination and visualization

The crystals were cryoprotected by supplementing the reservoir solution with 25% glycerol. Data collection was performed at the synchrotron beamline at Brookhaven National Laboratory (BNL), with datasets acquired over an oscillation range of 270° and an oscillation width of 0.2°. The crystal-to-detector distance was maintained at 200 mm throughout the data collection process. The collected datasets were processed using BNL’s inbuilt automated processing tools, and the integrated and scaled data were directly used for molecular replacement with PHASER (57). The initial model was generated using AlphaFold 2, and the atomic coordinates were refined against the electron density map in PHENIX (58), followed by structure building in Coot (59). Data collection and processing statistics are provided in Table S1. Figures were prepared using PyMOL (https://www.pymol.org/) and UCSF ChimeraX (60), with the final layout created in Adobe Photoshop.

## Supporting information

strains, plasmids and oligonucleotides

## ACKNOWLEDGMENTS

The authors thank all members of the Marraffini laboratory for helpful discussion and encouragement, B.R. Levin for providing the T4 phage, and A. Harms for providing the BASEL phage collection. C.F.B. was supported by an NIH T32 Chemistry-Biology Interface training grant (GM136640-Tan). Mass spectrometry data were generated by the Proteomics Resource Center at The Rockefeller University (RRID:SCR_017797) using instrumentation funded by the Sohn Conferences Foundation and the Leona M. and Harry B. Helmsley Charitable Trust. S.F.B. was funded in part by NIH R35GM122559. A.A.H. is currently a Jane Coffin Childs Memorial Foundation Postdoctoral Fellow at Memorial Sloan Kettering Cancer Center. D.J.P. is supported by the NIH (GM129430, AI141507, and GM145888), the Maloris Foundation, and a Memorial Sloan-Kettering core grant (P30-CA008748). This research used NSLS-II MX User Resources (FMX) of the National Synchrotron Light Source II, a US Department of Energy (DOE) Office of Science User Facility operated for the DOE Office of Science by Brookhaven National Laboratory under contract no. DE-SC0012704. The Center for BioMolecular Structure (CBMS) is primarily supported by the NIH National Institute of General Medical Sciences (NIGMS) through a Center Core P30 grant (P30GM133893) and by the DOE Office of Biological and Environmental Research (KP1605010). R.M. is supported by Austrian Science Fund FWF (grant no. F8011-B (10.55776/F80)). L.A.M. was supported by the Stavros Niarchos Foundation (SNF) as part of its grant to the SNF Institute for Global Infectious Disease Research at The Rockefeller University. L.A.M. is an investigator of the Howard Hughes Medical Institute.

## AUTHOR CONTRIBUTIONS

A.M.P., A.A.H. and L.A.M. designed and conceived the study. A.M.P. performed all in vivo and in vitro experiments with help from A.A.H. and C.F.B. Z.Z. performed all structural studies, including protein purification, X-ray crystallography and structural analyses, under the guidance of D.J.P. M.T. assisted with crystal picking and remote data collection. R.M. synthesized the oligonucleotides used in the crystallization of the proteins. C.P. and H.M. performed mass spectrometry. S.F.B. provided the soil metagenomic DNA libraries. K.B. prepared 5hmC-modified dsDNA under the supervision of R.M. A.M.P., Z.Z., A.A.H., R.M., D.J.P. and L.A.M. wrote and edited the manuscript.

## SUPPLEMENTARY FIGURES

**Figure S1.**
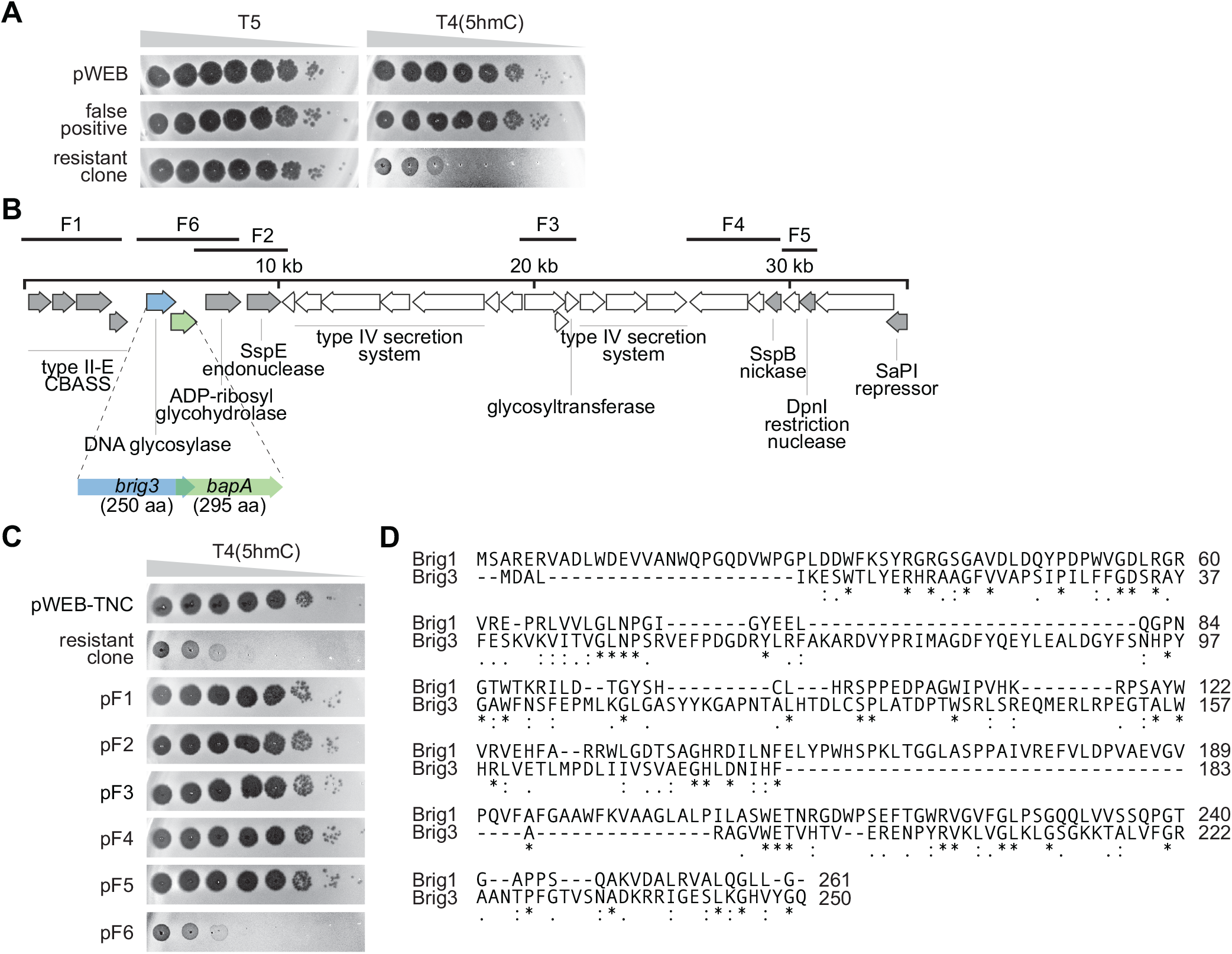
Isolation of Brig3 from an eDNA library. **(A)** Ten-fold serial dilutions of T5 or T4(5hmC) phage on lawns of *E. coli* EC100 cells generated from colonies sampled randomly from the eDNA library population that survived T4(5hmC) infection. Cells harboring an empty pWEB cosmid were used as a negative control. A representative result for a surviving colony that did not contain a *bona fide* immunity gene in the eDNA is shown. Plaquing on a lawn of cells that originated from a surviving colony that harbored a cosmid carrying immunity genes is also shown. **(B)** Genes present within the 35.4 kb soil metagenomic DNA fragment that provided T4 immunity. Subcloned regions (fragments F1-F6) are indicated. Putative defense genes are shown in grey. Genes labeled “*1*” and “*2*” correspond to *ORF1* (Brig3) and *ORF2* (BapA), respectively. **(C)** Ten-fold serial dilutions of phage T4 on lawns of *E. coli* EC100 carrying pWEB or cosmids containing fragments F1-F6. **(D)** Amino acid alignment of Brig1 and Brig3. (*), (:) and (.) mark identical, strongly conserved and weakly conserved residues, respectively.

**Figure S2.**
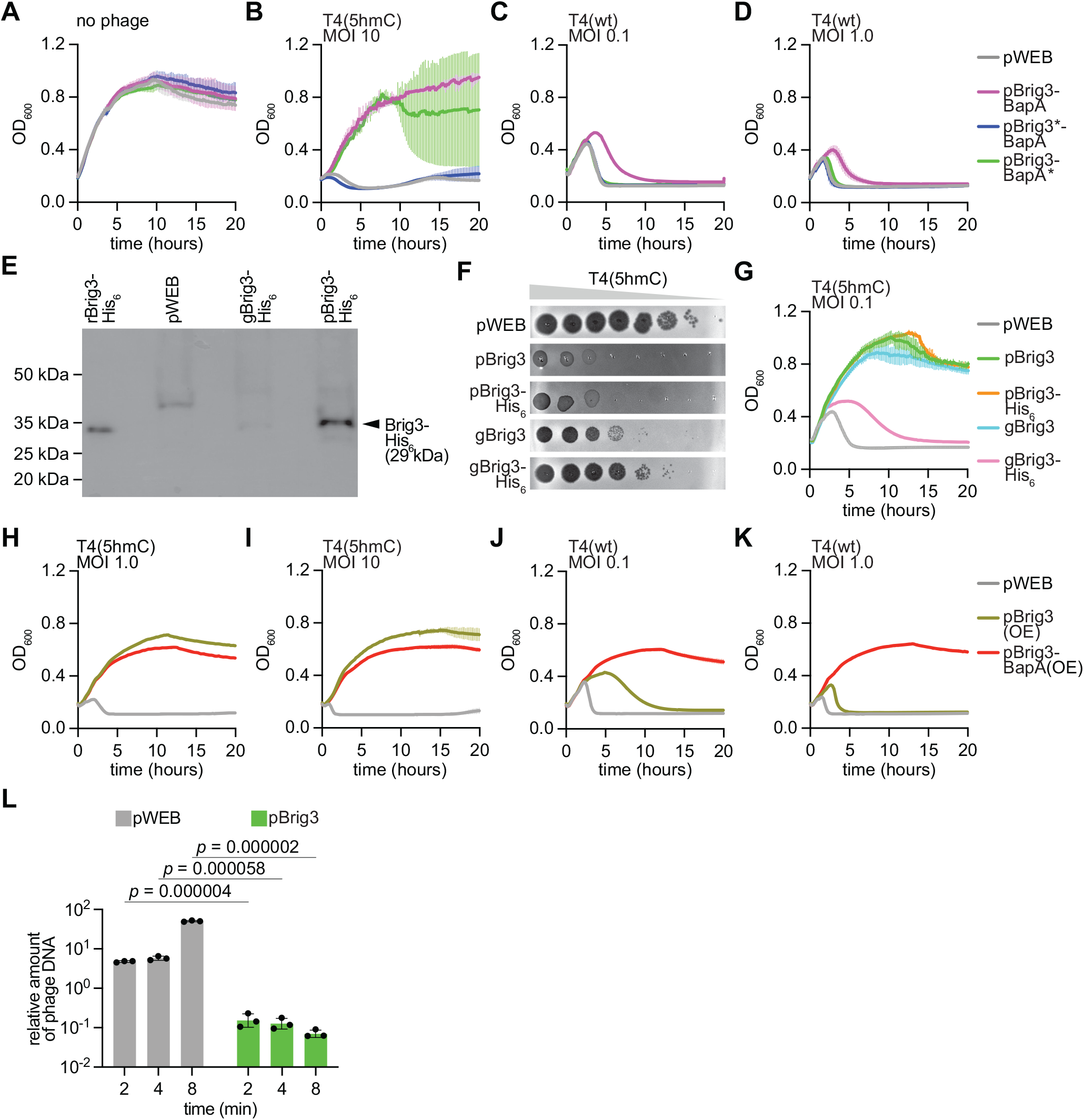
Infection of cultures expressing Brig3 and/or BapA. **(A-D)** Growth of *E. coli* EC100 that harbor the pWEB cosmid carrying wild-type and stop codon mutant versions of *ORF1* and/or *ORF2*, measured by optical density at 600 nm (OD_600_) either **(A)** uninfected, or after infection with **(B)** T4(5hmC) at MOI 10, **(C)** T4(wt) at MOI 0.1 or **(D)** T4(wt) at MOI 1.0. The mean ± S.D. for 3 biological replicates is shown. **(E)** Anti-His_6_ Western blot of *E. coli* EC100 lysates carrying the pWEB-TNC cosmid (pWEB), pBrig3-His_6_, or harboring a chromosomal copy of *brig3-His_6_* (gBrig3-His_6_). Recombinant Brig3-His_6_ (rBrig3-His_6_) was used as a positive control (4.5 ng of purified protein were loaded). The positions of molecular weight markers are displayed. **(F)** Ten-fold serial dilutions of T4(5hmC) phage on lawns of *E. coli* EC100 carrying the pWEB-TNC cosmid (pWEB), pBrig3, pBrig3-His_6_, or harboring a chromosomal copy of *brig3* or *brig3-His_6_* (gBrig3 and gBrig3-His_6_, respectively). **(G)** Growth of *E. coli* EC100 strains shown in (F) measured as OD_600_ after infection with T4(5hmC) phage at MOI 0.1. **(H-K)** Growth of *E. coli* EC100 that harbor the pWEB cosmid as a negative control or plasmids over-expressing either *ORF1* or *ORF1/ORF2*, measured by optical density at 600 nm (OD_600_) after infection with **(H)** T4(5hmC) at MOI 1.0, **(I)** T4(5hmC) at MOI 10, **(J)** T4(wt) at MOI 0.1 or **(K)** T4(wt) at MOI 1.0. The mean ± S.D. for 3 biological replicates is shown. **(L)** Quantitative PCR analysis of T4(5hmC) DNA through amplification of the *gp42* gene. Viral DNA was extracted from infected *E. coli* EC100 cells carrying pWEB or pBrig3 at 2, 4 and 8 minutes after the addition of phage at an MOI of 1. Fold-change values were calculated relative to the pWEB 2-minute time point. Mean ± SD values are reported for three independent experiments; *p*-values are reported for multiple unpaired Student T-tests.

**Figure S3.**
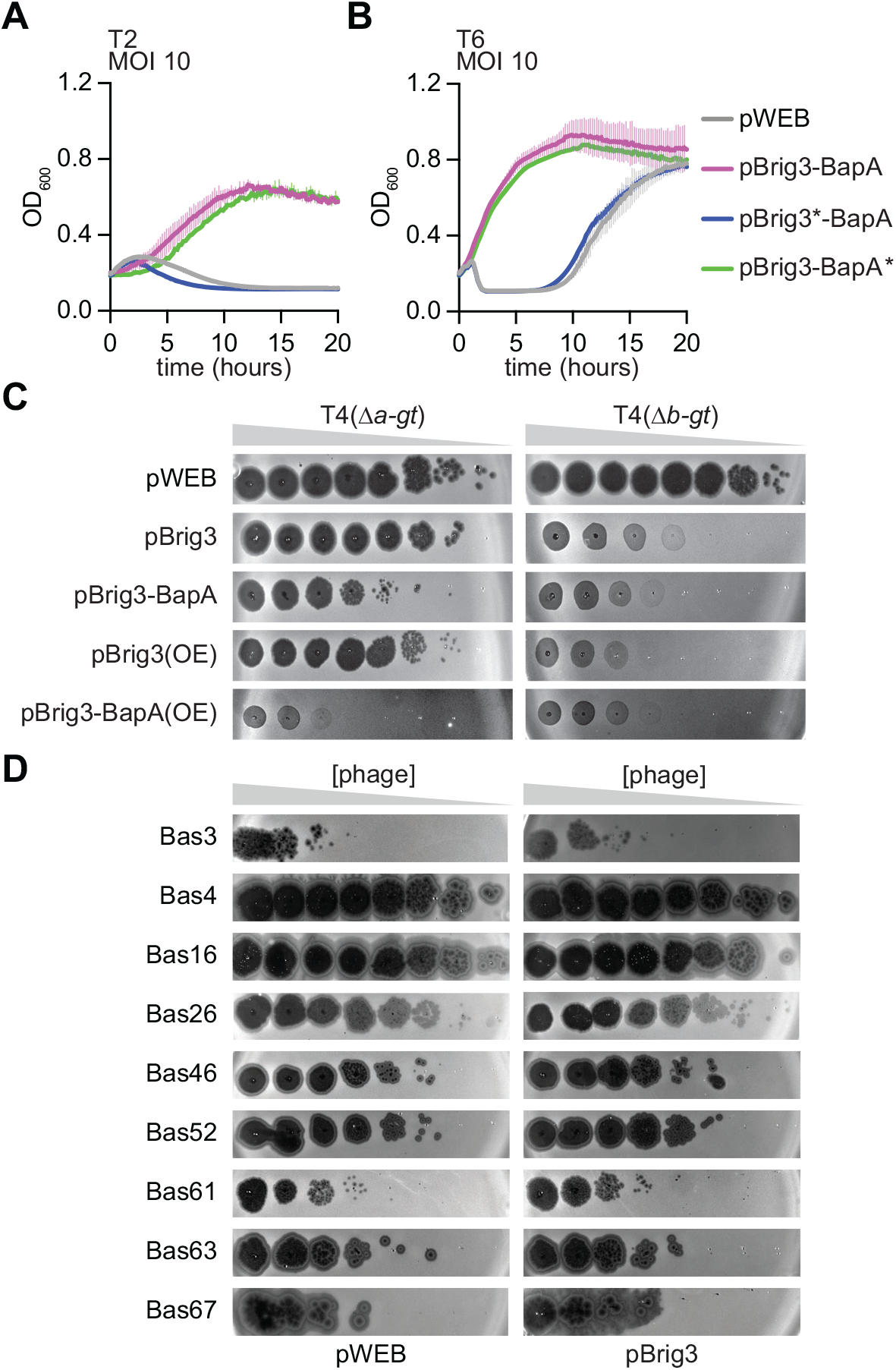
Brig3 immunity against diverse T-even phages. **(A)** Growth of *E. coli* EC100 that harbor the pWEB cosmid carrying wild-type and stop codon mutant versions of Brig3 and/or BapA, measured by optical density at 600 nm (OD_600_) after infection with T2 at MOI 10. **(B)** Same as **(A)** but after T6 infection. **(C)** Ten-fold serial dilutions of T4 mutants lacking either α- or β-glucosyl transferase, Δ*a-gt* or Δ*b-gt*, respectively, on lawns of *E. coli* EC100 that harbor the pWEB cosmid encoding Brig3 or Brig3-BapA, or a plasmid that over-expresses (OE) these proteins. **(D)** Ten-fold serial dilutions of non-*Tevenvirinae* phages from the Basel collection on lawns of *E. coli* EC100 that harbor the pWEB cosmid or pBrig3. Bas03, *Braunvirinae*; Bas04, *Tempevirinae*; Bas16, *Dhillonvirus*; Bas26, *Eseptimavirus*; Bas46, *Mosigvirus*; Bas52, *Vequintavirus*; Bas61, unclassified; Bas63, *Felixounavirus*; Bas67, *Berlinvirus*.

**Figure S4.**
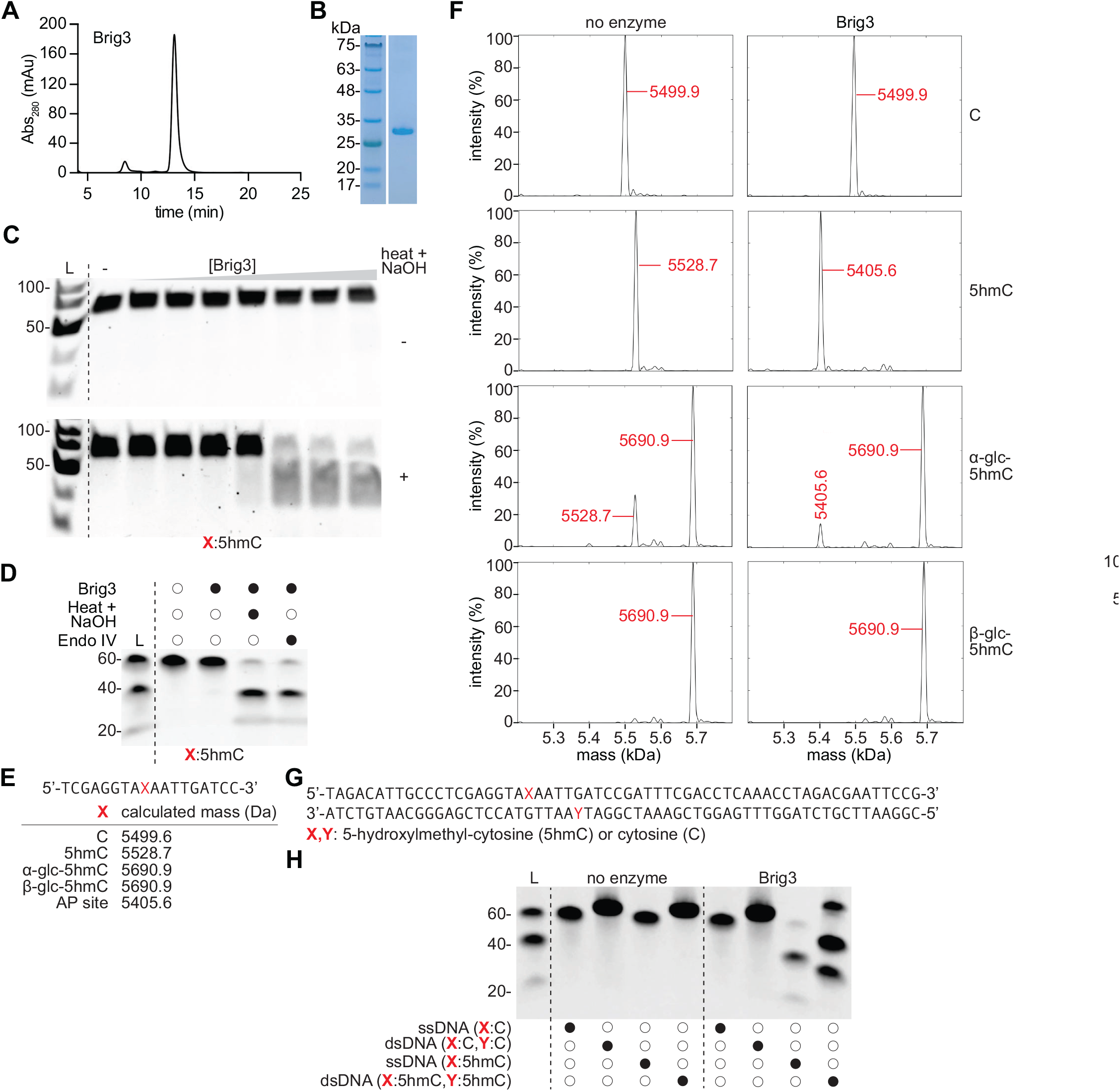
Brig3 introduces abasic sites in ssDNA and dsDNA oligonucleotides containing 5hmC. **(A)** Size-exclusion chromatography (SEC) analysis of Brig3-His_6_ purified through nickel affinity, using a Superdex 75 Increase column. **(B)** SDS-PAGE of the fraction corresponding to the SEC peak shown in **(A)**, visualized using Coomassie stain. **(C)** PAGE of 5hmC-containing oligonucleotides incubated either in the absence of Brig3 or with ten-fold dilutions of the enzyme Brig3 (10^-3^, 10^-2^, 10^-1^, 1, 10, 100, 1000 nM), at 37°C for 30 minutes, with and without heating in the presence of NaOH for 30 minutes. Gels were stained with ethidium bromide. L, ssDNA size ladder. **(D)** Same as **(C)** but including a reaction in which Brig3 treatment was followed by incubation with 50 units of Endonuclease IV at 37°C for 4 hours. **(E)** Calculated average masses, in daltons (Da), of the different 18-nucleotide oligonucleotides used for mass spectrometry. The central position (red “X”) is occupied by either cytosine (C), 5-hydroxymethylcytosine (5hmC) or α- or β-glucosyl-5hmC (α-glc-5hmC or β-glc-5hmC, respectively). **(F)** Deconvoluted zero-charge mass spectra from high resolution mass spectrometry of oligonucleotides harboring the nucleobases shown on the right at the central position, incubated with or without Brig3 at 37°C overnight. The masses of major peaks are indicated in red. **(G)** Sequence of complementary 60-nt single-stranded oligonucleotides used to test Brig3 activity on dsDNA. The red “X” and “Y” mark the positions occupied by modified 5hmC nucleobases. **(H)** PAGE of ssDNA and dsDNA oligonucleotides harboring different combinations of 5hmC or cytosines at “X” and “Y” positions, incubated with or without Brig3 at 37°C for 30 minutes, followed by heating in the presence of NaOH for 30 minutes. The gel was stained with ethidium bromide. L, ssDNA size ladder.

**Figure S5.**
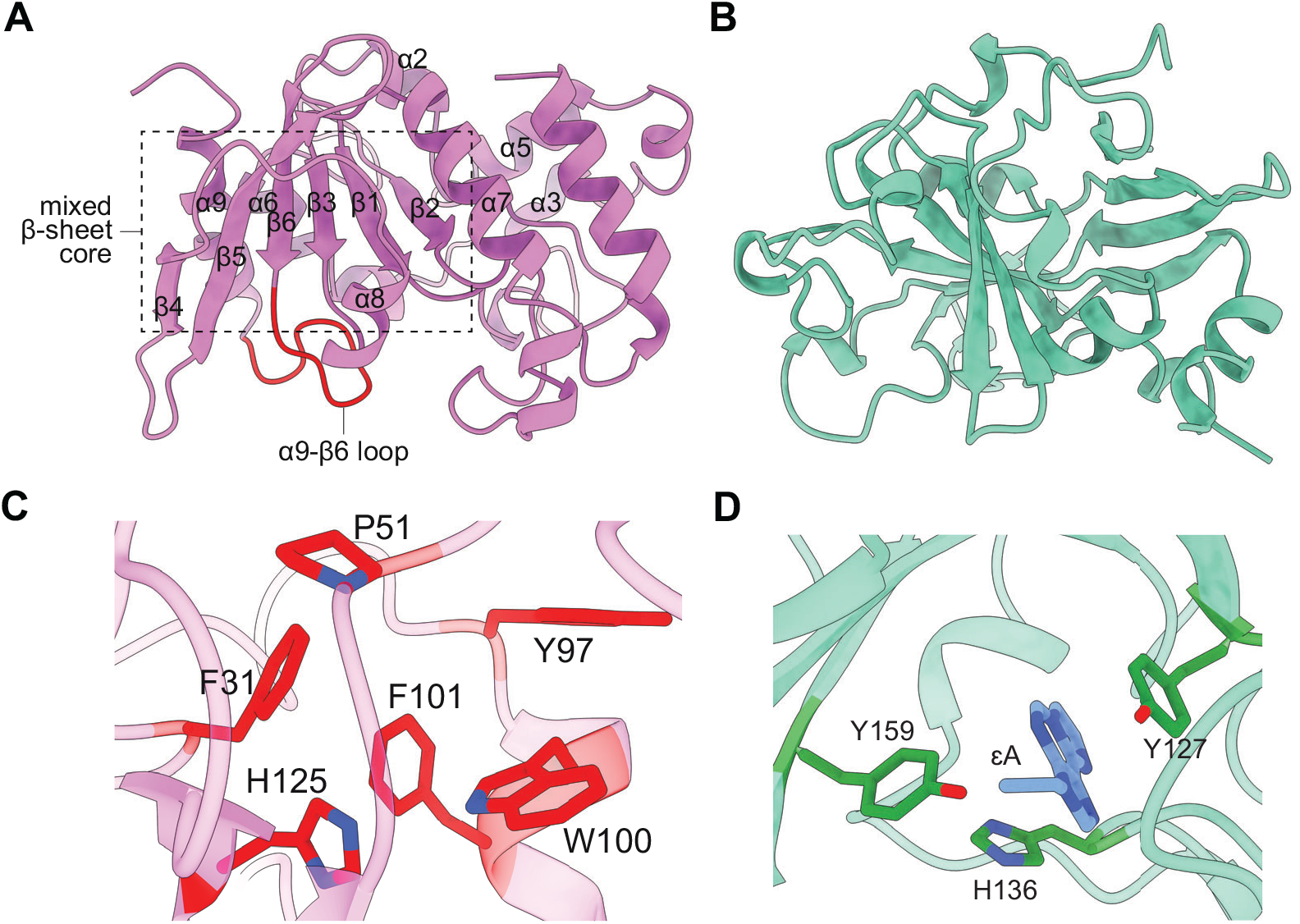
Crystal structure of wild-type apo-Brig3. **(A)** Crystal structure of wild-type Brig3, displaying a mixed α/β fold composed of nine α-helices and six β-strands. The curved, mixed β-sheet core and the α9-β6 insertion loop (in red) are labeled. **(B)** Crystal structure of alkyladenine DNA glycosylase (AAG), which also features a mixed α/β fold and shares a similar shape with Brig3 (PDB 1EWN). **(C)** Catalytic pocket of Brig3, lined with several aromatic residues. Key residues are labeled. **(D)** Active site of AAG, showing the flipped base ethenoadenine (ɛA) situated in the aromatic-lined pocket. Key residues are labeled.

**Figure S6.**
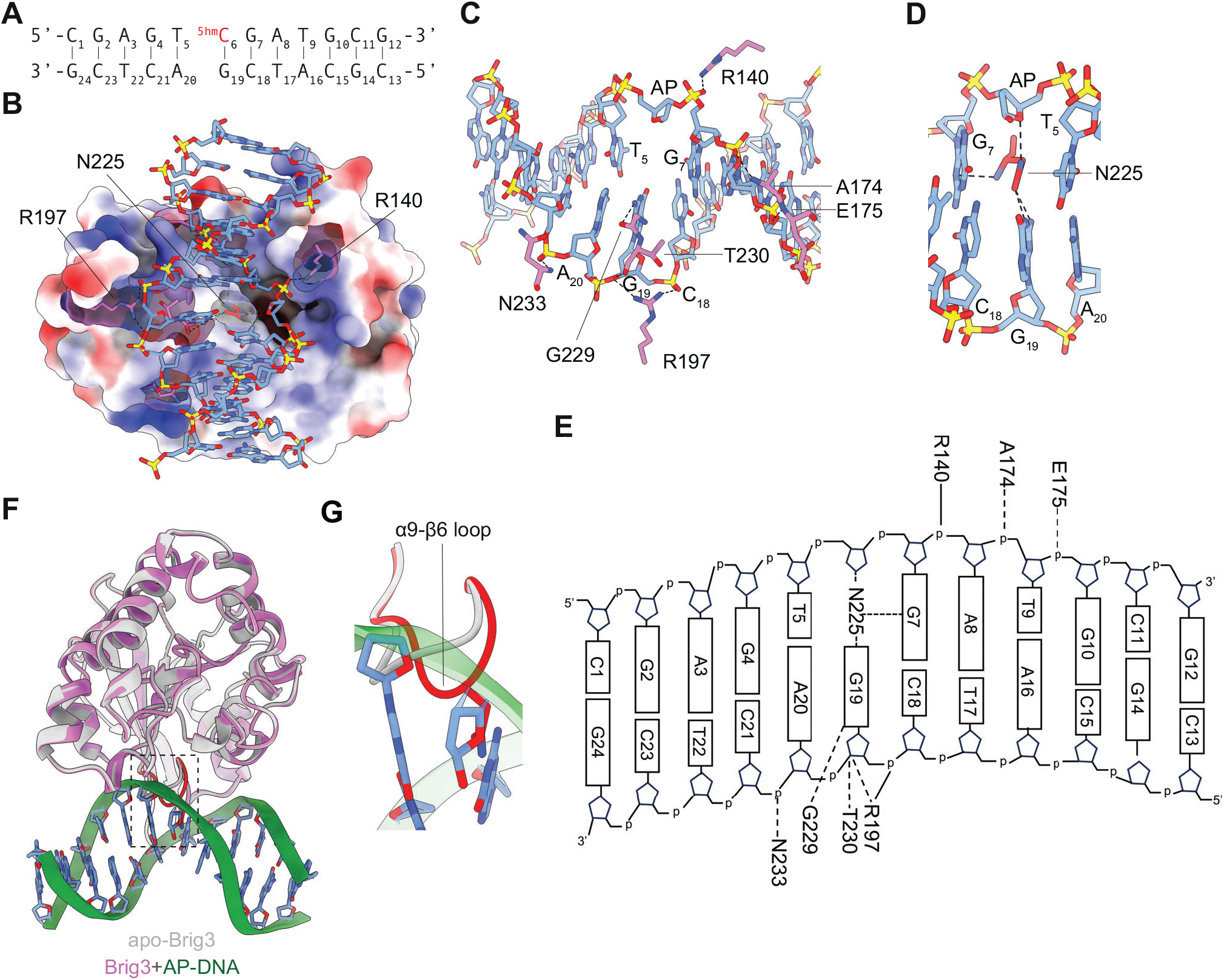
Crystal structure of wild-type Brig3 with a dsDNA substrate. **(A)** Sequence of the 12-mer dsDNA used for co-crystallization with Brig3, the 5-hydroxymethylcytosine (5hmC) nucleobase is shown in red. **(B)** Charge distribution map of the Brig3:dsDNA complex. Residues R197 and R140 interact with opposite DNA backbones, positioning the dsDNA substrate in a basic region of Brig3. N225 inserts into the abasic site and makes hydrogen bonds with the guanine that originally paired with 5hmC, an interaction that stabilizes the product of the base excision reaction. **(C)** Stable hydrogen bonds detected between residues R140, A174, E175, R197, G229, T230, and N233 with the DNA phosphate backbone and/or nucleobases surrounding the abasic site (AP). **(D)** Stable hydrogen bonds formed between residue N225, which is inserted into the abasic site (apurinic/apyrimidinic site, AP), and the adjacent (G_7_) and opposite (G_19_) nucleobases. **(E)** Schematic of specific interactions between Brig3 and the 12-mer dsDNA substrate shown in **(A)**. Solid and dotted lines indicate interactions via the amino acid side chain or the peptide backbone, respectively. **(F)** Structural alignment between the apo (silver) and dsDNA-bound (pink) forms of Brig3, with an RMSD of 0.361 Å for the protein components. The only notable difference shown in the boxed panel is found in the α9-β6 insertion loop, which is marked in red. **(G)** Region highlighted by the inset in **(F)** showing the most important difference in both structures: a shift in the α9-β6 insertion loop (displayed in red for the Brig3:dsDNA complex).

**Figure S7.**
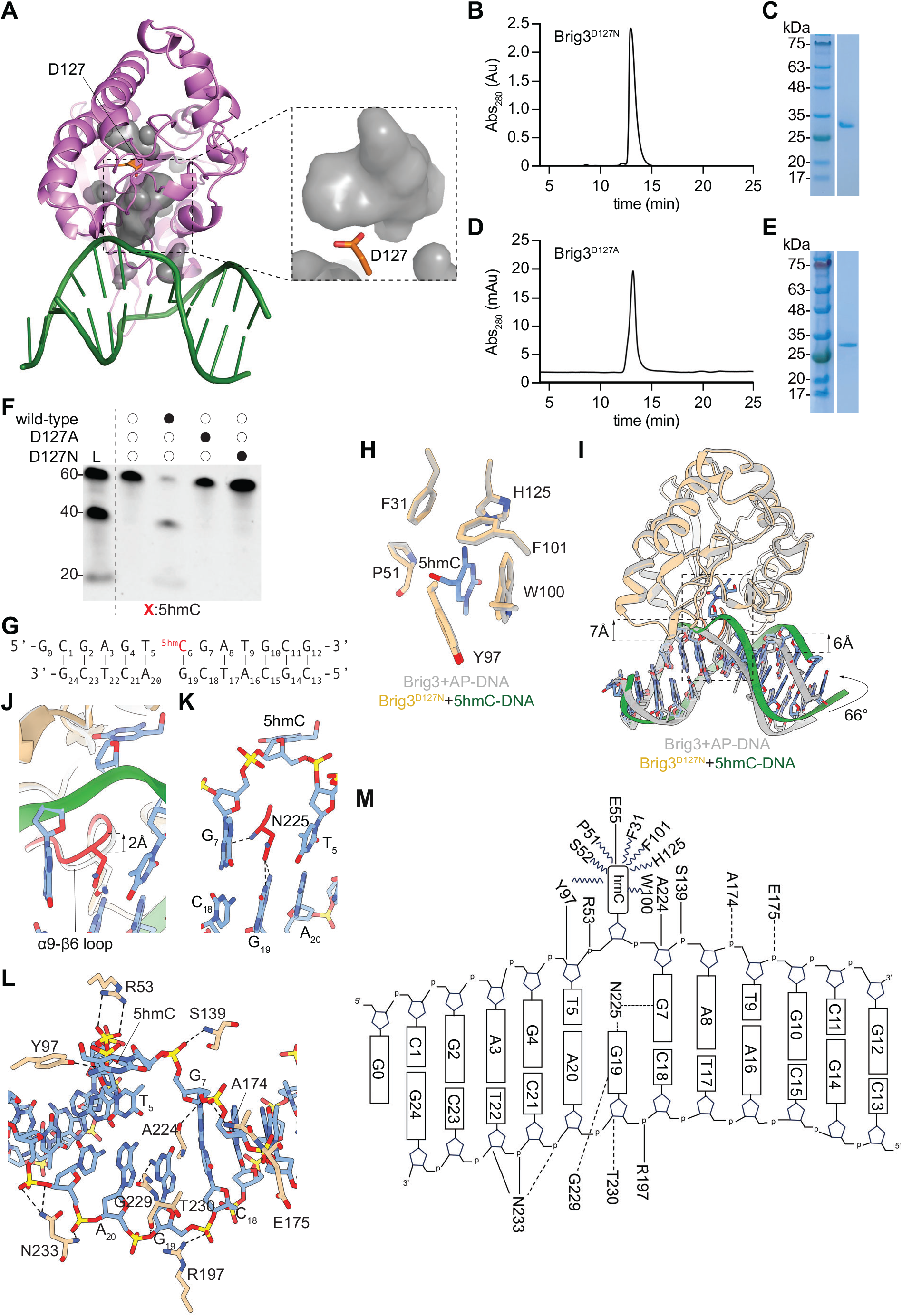
Crystal structure of Brig3^D127N^ with a dsDNA substrate. **(A)** Wild-type Brig3 (pink) bound to a 12-mer dsDNA duplex (green) with the structural cavities highlighted in translucent grey. Residue D127 is shown and colored in orange. The inset shows a zoomed view of the biggest cavity (presumably the active site) and its proximity to D127. **(B)** Size-exclusion chromatography (SEC) analysis of Brig3^D127N^-His_6_ purified through nickel affinity, using a Superdex 75 Increase column. **(C)** SDS-PAGE of the fraction corresponding to the SEC peak shown in **(B)**, visualized using Coomassie stain. **(D)** Same as **(B)** but for purified Brig3^D127A^-His_6_. **(E)** Same as **(C)** but for the Brig3^D127A^-His_6_ peak shown in **(D)**. **(F)** PAGE of 5hmC-containing oligonucleotides either untreated or incubated with wild-type and mutant (D127N, D127A) versions of Brig3 at 37°C for 30 minutes, followed by heating in the presence of NaOH for 30 minutes. Gels were stained with ethidium bromide. L, ssDNA size ladder. **(G)** Sequence of the 12-mer dsDNA used for co-crystallization with Brig3^D127N^, the 5-hydroxymethylcytosine (5hmC) nucleobase is shown in red. **(H)** Structural alignment of the catalytic pocket in wild-type (silver) and D127N (gold) Brig3, both in complex with a dsDNA substrate harboring an abasic site (apurinic/apyrimidinic site, AP) or a 5hmC nucleobase (in blue), respectively. **(I)** Structural alignment of wild-type (silver) and D127N (gold) Brig3, both in complex with a dsDNA substrate harboring an abasic site (AP, in silver) or a 5hmC nucleobase (in green), respectively, showing an RMSD of 0.396 Å. Protein structures exhibit minimal conformational changes, with the α9-β6 insertion loop showing the only noticeable shift. The changes in the dsDNA substrate include a shift of approximately 7 Å and 6 Å for the 5hmC-containing and complementary strands, as well as bending of 66° towards the Brig3^D127N^ protein. **(J)** Region highlighted by the inset in **(I)**, showing the change in the α9-β6 insertion loop of approximately 2 Å toward the phosphate backbone. **(K)** Stable hydrogen bonds formed between residue N225, which is inserted in place of the flipped 5hmC nucleobase, and the adjacent (G_7_) and opposite (G_19_) nucleobases. **(L)** Stable hydrogen bonds detected between residues R53, Y97, S139, A174, E175, R197, A224, G229, T230, and N233 with the DNA phosphate backbone and/or nucleobases surrounding the 5hmC nucleobase. **(M)** Schematic of specific interactions between Brig3^D127N^ and the 12-mer dsDNA substrate shown in **(G)**. Solid and dotted lines indicate interactions via the amino acid side chain or the peptide backbone, respectively.

**Figure S8.**
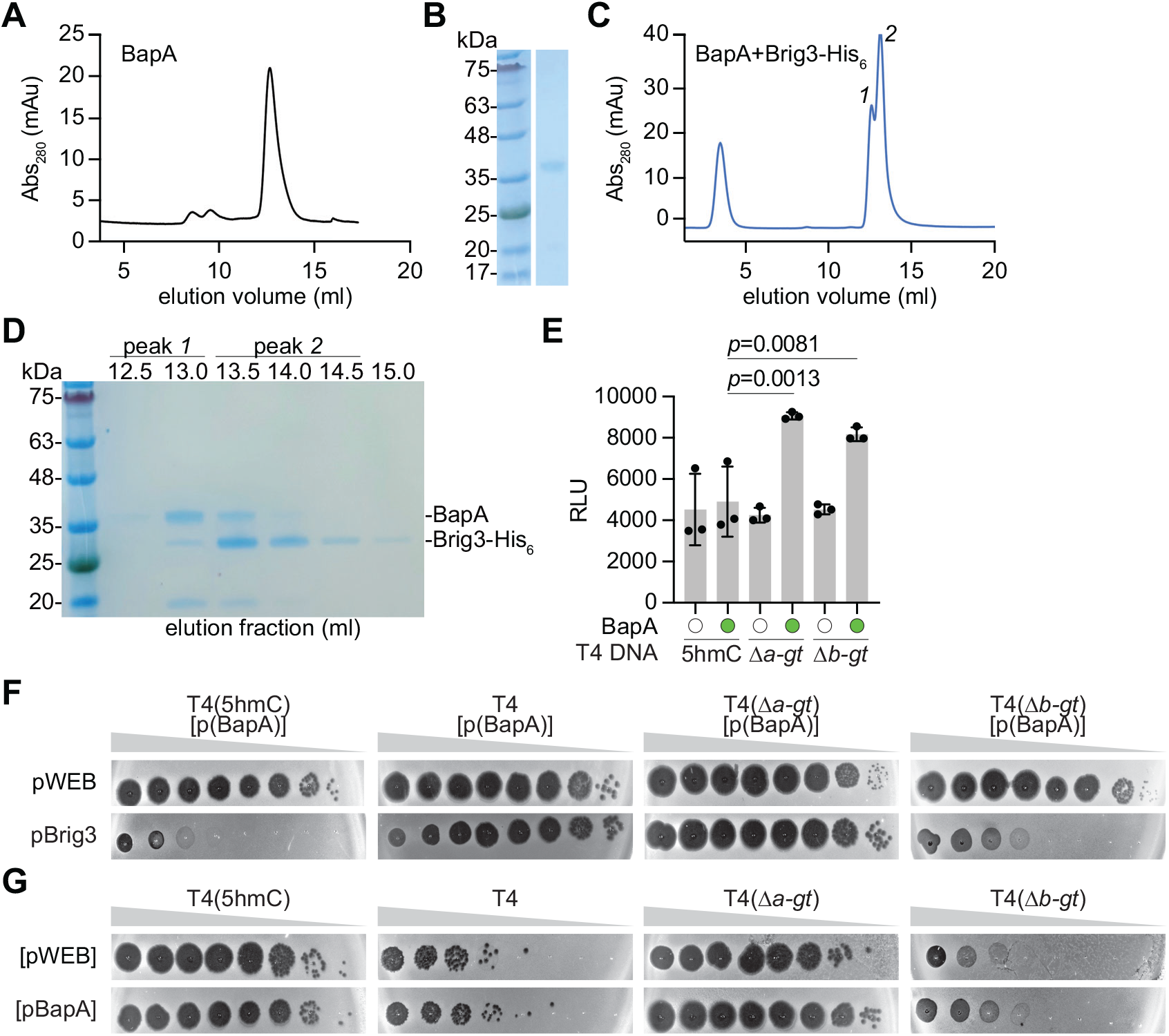
Purification and activity of BapA. **(A)** Size-exclusion chromatography (SEC) analysis of BapA after purification through nickel affinity and cleavage of the His_6_-SUMO tag, using a Superdex 75 Increase column. **(B)** SDS-PAGE of the fraction corresponding to the SEC peak shown in **(A)**, visualized using Coomassie stain. **(C)** In vitro assembly of BapA and Brig3-His_6_ at a 1:1.5 molar ratio, incubated on ice for one hour, and applied to a Superdex 75 Increase 10/300 column. The two distinct peaks are labeled “*1*” and “*2*”. **(D)** SDS-PAGE analysis of the SEC fractions encompassing both peaks. The separate elution profiles suggest that the two proteins do not form a stable complex under these in vitro assembly conditions. **(E)** Luminescence assay for glucose detection after incubation of genomic DNA extracted from different T4 phages with BapA overnight at 37°C. RLU, relative luminescence units. Error bars represent the standard error of the mean; *p*-values are reported for a Ordinary One-Way ANOVA with multiple comparisons. **(F)** Ten-fold serial dilutions of different T4 phage stocks obtained after amplification in *E. coli* EC100(pBapA) hosts (“[pBapA]”), plated on lawns of *E. coli* EC100 that harbor the pWEB or pBrig3 cosmids. **(G)** Ten-fold serial dilutions of different T4 phage stocks obtained after amplification in *E. coli* EC100 hosts harboring either pBapA (“[pBapA]”) or a control cosmid ([pWEB]), plated on lawns of *E. coli* EC100 that express Brig1.

**Figure S9.**
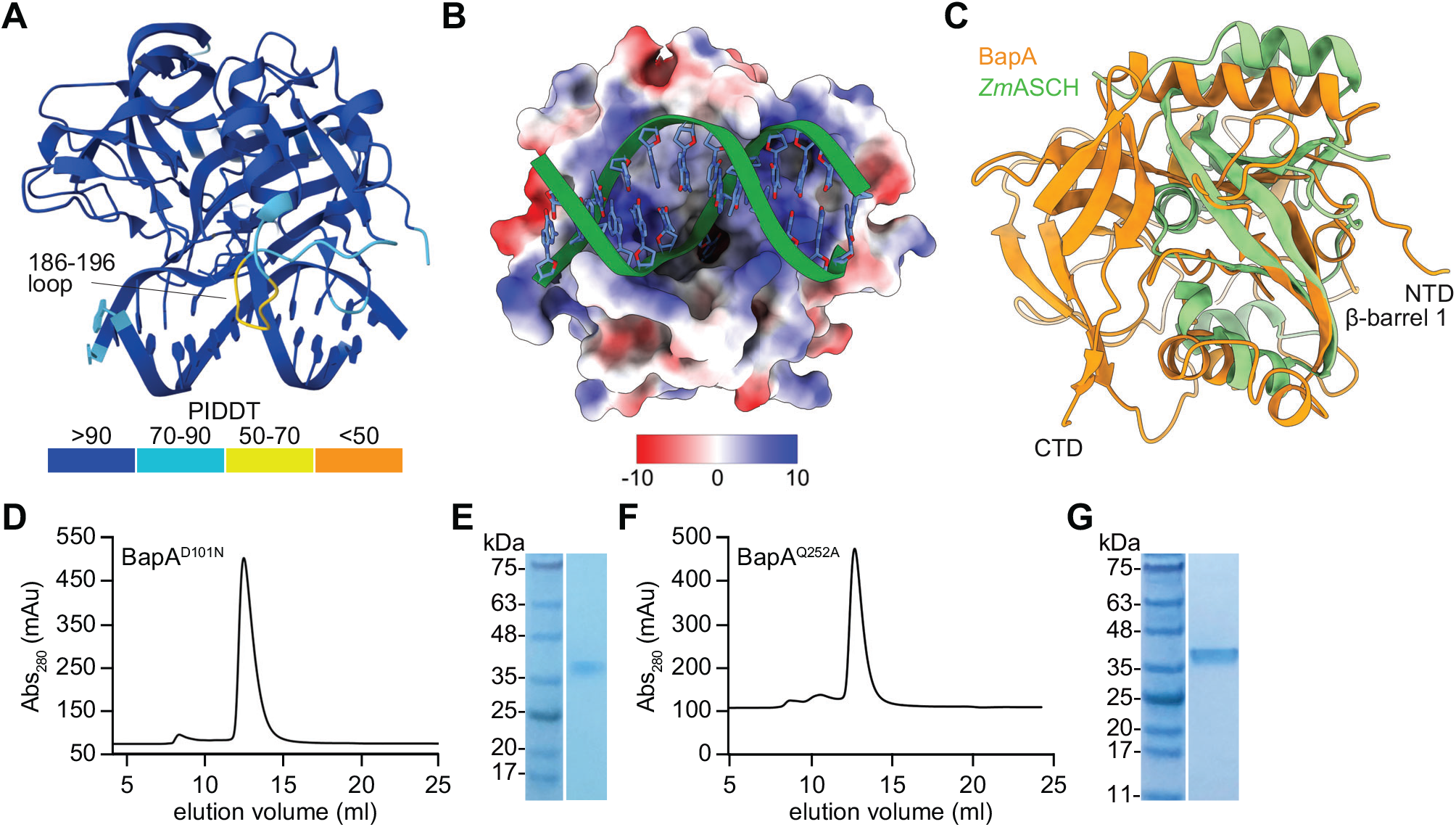
BapA structural model and purification of mutants. **(A)** AlphaFold 3 structural model of BapA in complex with a dsDNA substrate that harbors a 5hmC nucleobase, colored according with pIDDT confidence. The flexible loop 186-196 shows the lowest prediction confidence, consistent with its expected structural flexibility. **(B)** Electrostatic surface representation of the predicted BapA:dsDNA complex, showing the DNA duplex accommodated within a positively charged channel. **(C)** Structural comparison of BapA (orange) with an ASCH domain-containing protein from *Zymomonas mobilis*, *Zm*ASCH (green; PDB 5GUQ) showing a conserved β-barrel 1 topology (RMSD 1.108 Å). **(D)** Size-exclusion chromatography (SEC) analysis of BapA^D101N^ after purification through nickel affinity and cleavage of the His_6_-SUMO tag, using a Superdex 75 Increase column. **(E)** SDS-PAGE of the fraction corresponding to the SEC peak shown in **(D)**, visualized using Coomassie stain. **(F)** Same as **(D)** but for purified BapA^Q252A^. **(G)** Same as **(E)** but for the BapA^Q252A^ peak shown in **(F)**.

**Figure S10.**
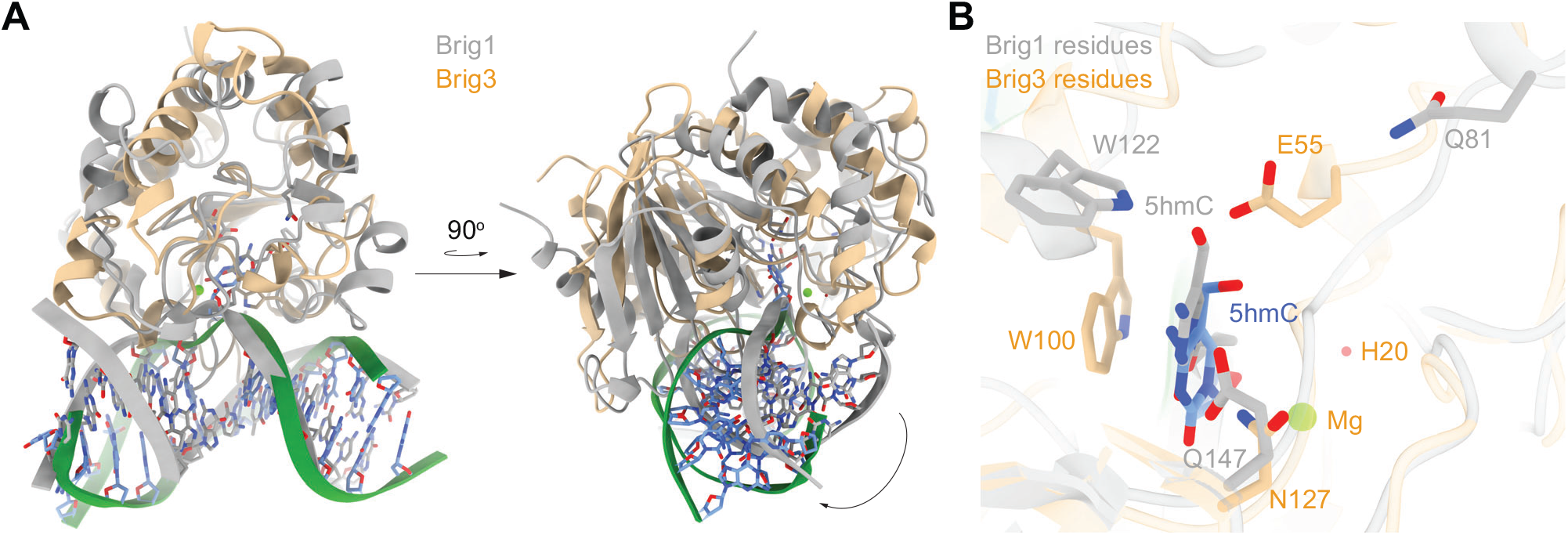
Structural comparison of Brig1:dsDNA and Brig3^D127N^:dsDNA complexes. **(A)** Overlay of an AlphaFold 3-generated Brig1:dsDNA complex (silver; ipTM = 0.91, pTM = 0.93, the dsDNA contains a single 5hmC nucleobase) with the crystal structure of Brig3^D127N^:dsDNA (gold). Structures are shown in two orientations. Both structures position the dsDNA duplex within a similar cleft, yielding an overall RMSD of 1.075 Å between the protein structures. The similarities in the geometry of DNA docking suggests a conserved mode of substrate recognition across members of the Brig family. The arrow indicates the shift in the DNA backbone caused by conformational changes in the three N-terminal α-helices of Brig1 and Brig3^D127N^. **(B)** Detailed view of the catalytic pockets within the Brig1:dsDNA model (silver) and the Brig3^D127N^:dsDNA (gold) crystal structure, highlighting residues involved in 5hmC recognition and catalysis. The 5hmC base is coordinated by Q81, W122, and Q147 in Brig1, and by E55, W100, and N127 in Brig3^D127N^, which also employs a Mg²⁺ ion and a coordinated water molecule. Brig1 exhibits a larger, more open pocket that extends further from the DNA, suggesting a potentially expanded catalytic space for accommodating the α-glc-5hmC substrate.

**Figure S11.**
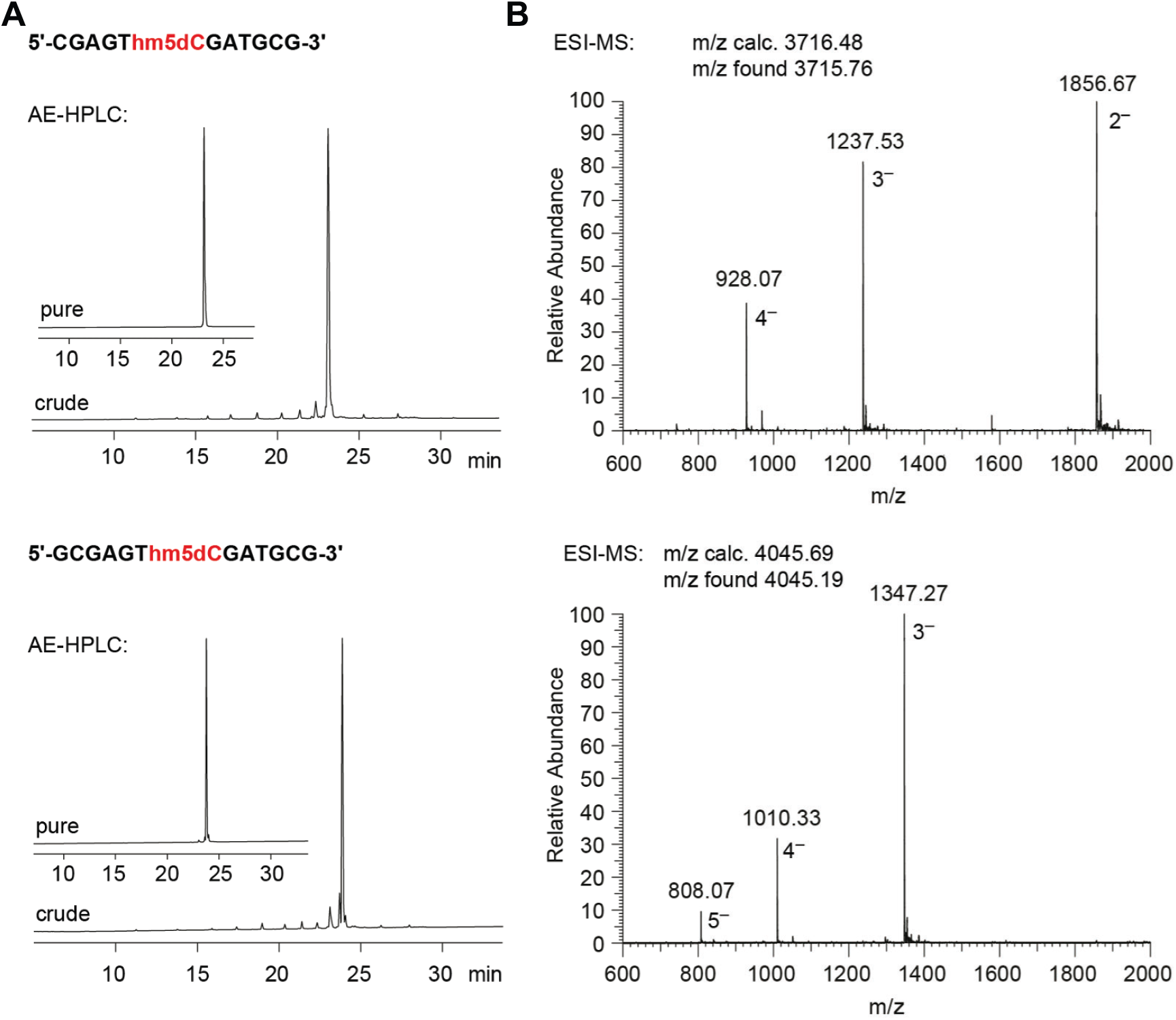
Characterization data of synthetic hm5dC-modified DNA oligonucleotides. **(A)** AE-HPLC Dionex DNApac was performed using a column, 80 °C, 1 mL/min, 0-40% of buffer B within 30 min, Buffer A: Tris-HCl (25 mM) and NaClO4 (20 mM) in aq 20% ACN, pH 8.0; Buffer B: Tris-HCl (25 mM) and NaClO_4_ (0.6 M) in aq 20% acetonitrile, pH 8.0. **(B)** LC-ESI MS was performed on Finnigan LCQ Advantage Max in the negative-mode (C18 XBridge column (2.5 µm, 2.1 mm × 50 mm), flow rate of 0.1 mL/min, gradient 0 to 100% B at 30 °C (eluent A: 8.6 mM triethylamine, 100 mM 1,1,3,3,3-hexafluoroisopropanol in H_2_O; eluent B: methanol).

