## Supplementary material for "A base-modifying enzyme extends the range of anti-phage defense provided by DNA glycosylases": Table S1

### Table S1. Data collection and refinement statistics (molecular replacement)

| | Brig3 apo | Brig3+12-mer dsDNA | Brig3 (D127N) +5'-overhang-containing 12-mer dsDNA |
|---|---|---|---|
| **Data collection** | | | |
| Space group | $P12_11$ | $P2_12_12_1$ | $C222_1$ |
| Cell dimensions | | | |
| *a*, *b*, *c* (Å) | 97.323, 95.339, 110.043 | 65.885, 73.479, 115.689 | 90.8, 96.839, 79.173 |
| $\alpha, \beta, \gamma$ (°) | 90, 100.909, 90 | 90, 90, 90 | 90, 90, 90 |
| Resolution (Å) | 47.67-2.894 (2.998-2.894) | 45.45-1.76 (1.823-1.76) | 33.12-1.61 (1.668-1.61) |
| $R_{sym}$ or $R_{merge}$ | 9.3e-18 (1.249e-17) | 0.02845 (0.8354) | 0.06337 (1.112) |
| $I / \sigma I$ | 7.85 (2.59) | 11.51 (0.84) | 5.02 (0.53) |
| Completeness (%) | 97.05 (90.66) | 99.68 (88.78) | 99.33 (88.90) |
| Redundancy | 1.0 (1.0) | 2.0 (2.0) | 2.0 (2.0) |
| **Refinement** | | | |
| Resolution (Å) | 2.9 | 1.8 | 1.6 |
| No. reflections | 42963 (3950) | 99986 (11097) | 40417 (4451) |
| $R_{work}$ / $R_{free}$ | 0.2063/0.2854 | 0.213/0.243 | 0.207/0.239 |
| No. atoms | 15909 | 4655 | 2694 |
| Macromolecules | 15909 | 4420 | 2508 |
| Ligand/ion | 0 | 12 | 1 |
| Water | 0 | 223 | 185 |
| *B*-factors | 39.47 | 37.44 | 36.49 |
| Macromolecules | 39.47 | 36.10 | 28.01 |
| Ligand/ion | 0 | 29.89 | 39.32 |
| Water | 0 | 38.09 | 35.41 |
| R.m.s. deviations | | | |
| Bond lengths (Å) | 0.01 | 0.008 | 0.013 |
| Bond angles (°) | 1.24 | 1.12 | 1.47 |

*Number of xtals for each structure: One crystal. *Values in parentheses are for highest-resolution shell.
